# Ocean acidification induces subtle shifts in gene expression and DNA methylation in mantle tissue of the Eastern oyster (*Crassostrea virginica*)

**DOI:** 10.1101/2020.06.05.136424

**Authors:** Alan M. Downey-Wall, Louise P. Cameron, Brett M. Ford, Elise M. McNally, Yaamini R. Venkataraman, Steven B. Roberts, Justin B. Ries, Katie E. Lotterhos

## Abstract

Early evidence suggests that DNA methylation can mediate phenotypic responses of marine calcifying species to ocean acidification (OA). Few studies, however, have explicitly studied DNA methylation in calcifying tissues through time. Here, we examined the phenotypic and molecular responses in the extrapallial fluid and mantle (fluid and tissue at the calcification site) in the Eastern oyster (*Crassostrea virginica*) exposed to experimental OA over 80 days. Oysters were reared under three experimental pCO_2_ treatments (‘control’, 580 μatm; ‘moderate OA’, 1000 uatm; ‘high OA’, 2800 μatm) and sampled at 6 time points (24 hours - 80 days). We found that high OA initially induced changes in the pH of the extrapallial fluid (pH_EPF_) relative to the external seawater, but the magnitude of this difference was highest at 9 days and diminished over time. Calcification rates were significantly lower in the high OA treatment compared to the other treatments. To explore how oysters regulate their extrapallial fluid, gene expression and DNA methylation were examined in the mantle-edge tissue of oysters from day 9 and 80 in the control and high OA treatments. Mantle tissue mounted a significant global molecular response (both in the transcriptome and methylome) to OA that shifted through time. Although we did not find individual genes that were significantly differentially expressed to OA, the pH_EPF_ was correlated with the eigengene expression of several co-expressed gene clusters. A small number of OA-induced differentially methylated loci were discovered, which corresponded with a weak association between OA-induced changes in genome-wide gene body DNA methylation and gene expression. Gene body methylation, however, was not significantly correlated with the eigengene expression of pH_EPF_ correlated gene clusters. These results suggest that in *C. virginica*, OA induces a subtle response in a large number of genes, but also indicates that plasticity at the molecular level may be limited. Our study highlights the need to re-assess the plasticity of tissue-specific molecular responses in marine calcifiers, as well as the role of DNA methylation and gene expression in mediating physiological and biomineralization responses to OA.

## Introduction

Ocean acidification (OA), a decrease in seawater pH due to the uptake of anthropogenic CO_2_, is expected to have substantial effects on marine species and ecosystems in the near future (Orr et al., 2005; Guinotte and Fabry, 2008; Doney et al., 2009). Specifically, OA is driving a shift in the carbonate system equilibrium, resulting in decreased availability of carbonate ions and lower calcium carbonate saturation state (Feely et al., 2004; Orr et al., 2005). This may be particularly problematic for marine calcifying species, which build their shells and skeletons from calcium and carbonate ions.

Substantial effort has been invested to evaluate the short and long-term consequences of OA across taxa and life history stages. Generally, prolonged exposure to OA tends to have negative effects on calcification, metabolism, and growth, as observed in corals (e.g., Anthony et al., 2008), pteropods, gastropods (e.g., Melatunan et al., 2013), and bivalves (e.g., Talmage and Gobler, 2010). However, these negative effects are not universal, varying in direction (negative or positive) and intensity depending on taxon, severity of OA, and duration of exposure (Ries et al., 2009; Kroeker et al., 2010). Observed non-negative effects may be the result of individual resilience or the capacity of certain species to acclimatize to OA. One way this may occur is via the maintenance of pH homeostasis at the site of calcification despite an increasingly acidic environment, which is a polyphyletic response to ocean acidification (Liu et al., 2018) that has been observed in scleractinian corals (Al-Horani et al., 2002; Ries, 2011; Venn et al., 2011; McCulloch et al., 2012; Holcomb et al., 2014), foraminifera (Rink et al., 1998; Köhler-Rink and Kühl, 2000; de Nooijer et al., 2008, 2009), calcareous green algae (De Beer and Larkum, 2001), coralline red algae (Donald et al., 2017; Anagnostou et al., 2019; Liu et al., 2020), coccolithophores (Liu et al. 2018), and bivalves (Ramesh et al., 2017; Cameron et al., 2019). In bivalves, calcification occurs within the extrapallial fluid (EPF) located between the shell and mantle epithelium, the composition of which is regulated via the active exchange of ions and other constituents through the mantle epithelium (Crenshaw and Neff, 1969; Crenshaw, 1972). However, we are just beginning to investigate the specific genes or molecular mechanisms within the mantle tissue that regulate this process (Rajan and Vengatesen, 2020), and the potential trade-offs that exist for maintaining EPF chemistry that is supportive of calcification in the face of long-term OA exposure.

Transcriptomic studies have proven to be a powerful approach for identifying genes and understanding pathways that shape organismal response to OA, and have been used to help elucidate potential trade-offs that result from acclimatization (Evans and Hofmann, 2012), including genes associated with biomineralization, acid-base regulation, and metabolic function (Evans et al., 2013; Davies et al., 2016; Li et al., 2016; Goncalves et al., 2017; Wong et al., 2018; Griffiths et al., 2019). From this literature it has been hypothesized that there is a trade-off between maintaining calcification versus other core functions (Wood et al., 2008), whereby increased or sustained calcification under continued exposure may become too costly over time. At the transcriptomic level, this cost can manifest as shifts in the expression of both calcification and metabolic genes through time (Li et al., 2016), suggesting that certain acclimatization responses may be unsustainable during extended exposure to OA. The precise mechanisms that mediate these changes in gene expression remain underexplored, but recent studies indicate that DNA methylation, an epigenetic modification, may be an important regulator of these responses (Li et al., 2018; Liew et al., 2018; Bogan et al., 2020).

DNA methylation is an important mediator of gene regulation and physiological response across a diverse range of taxa (see reviews Zilberman et al., 2007; Zemach et al., 2010; Schübeler, 2015; Zhang et al., 2018). DNA methylation refers to the attachment of a methyl group to cytosine, and is most frequently studied in the context of the cytosine-guanine (CpG) motif. These modifications occur on top of the genome without altering the underlying DNA sequence (Richards, 2006; Bonduriansky and Day, 2018), and may be stable enough to be passed from parent to offspring (Jablonka and Lamb, 2002). Previous work has shown they can have an important role in determining an organism’s phenotype across a range of species, including fur color and obesity in mice (Dolinoy et al., 2006), rat pup behavior as determined by parental care (Anier et al., 2014), response to phosphate starvation in *Arabidopsis* (Yong-Villalobos et al., 2015), and caste phenotypes in social insects (Lyko et al., 2010). Importantly, the precise manner in which DNA methylation serves to regulate phenotype is taxon-specific.

In invertebrates, correlative studies have begun to elucidate the relationship between DNA methylation, gene expression, and phenotype (Feil and Fraga, 2012; Roberts and Gavery, 2012; Metzger and Schulte, 2016). In basal invertebrates (e.g., corals and oysters), genomes are typically only sparsely methylated compared to vertebrates, and DNA methylation tends to be concentrated in CpGs within gene bodies (i.e., exons and introns). In this group, gene body methylation was found to be positively correlated with gene expression and negatively correlated with the variation in expression among individuals (Gavery and Roberts, 2013; Liew et al., 2018). Based on these patterns, DNA methylation may be acting to regulate not only the level of expression, but also the amount of transcriptional noise among individuals (Huh et al., 2013). Furthermore, recent evidence suggests that unlike vertebrates, non-deuterostome invertebrate DNA methylation is not necessarily erased during embryogenesis, opening up the possibility for transmission of marks between generations (Xu et al., 2019).

Understanding the role of environment induced DNA methylation in regulating gene expression and phenotypic plasticity is of particular interest in the context of marine systems and global change (see reviews, Hofmann, 2017; Eirin-Lopez and Putnam, 2019). In corals, DNA methylation was shown to respond to environmental change and was correlated with gene expression (Dixon et al., 2018). OA-induced changes in DNA methylation have also been observed in some species of coral (Putnam et al., 2016; Liew et al., 2018), suggesting a potential regulatory role of DNA methylation in coral response to OA. Liew et al. (2018) found OA-sensitive DNA methylation also corresponded with biomineralization-related traits, supporting the idea that DNA methylation may have an important role in phenotypic plasticity and acclimatization to OA in corals. Although these studies examined OA reponses in many different tissues, none of these studies quantified methylation responses to OA at base-pair resolution in calcifying tissue.

Here, we examine the responses of the Eastern oyster *(Crassostrea virginica)* to OA in the tissue (mantle) and extrapallial fluid (EPF) that are involved in oyster calcification. *Crassostrea virginica* is a coastal marine bivalve that provides critical ecosystem services in the form of habitat structure and water filtration, and also serves as an economically important food source (Ekstrom et al., 2015; Gómez-Chiarri et al., 2015). Its genome has been recently sequenced (Gómez-Chiarri et al., 2015) and the species has been the subject of extensive OA studies. Consistent with other bivalve species, *C. virginica* exhibits largely negative calcification and physiological responses to OA (Ries et al., 2009; Beniash et al., 2010; Waldbusser et al., 2011; Ivanina et al., 2013; Matoo et al., 2013; Gobler and Talmage, 2014), with larvae appearing particularly susceptible (Miller et al., 2009; Talmage and Gobler, 2009) and juveniles exhibiting relative resilience (Dodd et al., 2015). Measurements of oyster EPF chemistry suggests that they, like other bivalve species, promote calcification by regulating the chemical composition of the EPF via the active exchange of ions and other constituents through the mantle epithelium (Crenshaw and Neff, 1969; Crenshaw, 1972). Estimates of bivalve pH_EPF_ obtained from proton-sensitive microelectrodes and shell boron-isotope chemistry suggest that some bivalve species regulate the carbonate chemistry of this fluid under OA conditions (Cameron et al., 2019; Liu et al., 2020). However, it is unclear if this response is sustainable under long-term OA exposure and little is known about the genes that underpin these phenotypic responses, or the role of DNA methylation as a potential mediator of these processes.

The primary goal of this study was to investigate the molecular mechanisms underlying the biomineralization response of *C. virginica* to short- and long-term OA exposure. Specifically, the role of DNA methylation in physiological and gene expression responses in *C. virginica* mantle-edge tissue. We did this through a controlled laboratory experiment in which oysters were exposed to three levels of CO_2_-induced OA (control, moderate, and high) and monitored for calcification rate and pH_EPF_ over 80 days. Based on patterns we observed in the pH_EPF_ response, we examined gene expression and DNA methylation responses in mantle-edge tissue at two time points (day 9 and 80) for oysters exposed to low and high OA treatments. We hypothesize that the oysters mitigate the effects of OA by elevating pH_EPF_ relative to seawater pH and that this process is associated with a molecular response in the oyster transcriptome and methylome. By establishing the first integrated dataset on EPF carbonate chemistry, gene expression, and DNA methylation in a mollusk exposed to variable degrees of ocean acidification, our study provides new insights into the responses of bivalves to OA over short and long timescales.

## Materials and methods

### Oyster collection and preparation

Adult *C. virginica* were collected from three intertidal sites within Plum Island Sound, Massachusetts, USA (Site 1, 42.751636, −70.837023; Site 2, 42.725186, −70.855022; Site 3, 42.681764, −70.813498) in late April 2017. These sites are all within 8 km of each other and did not represent distinct genetic populations. However, to compensate for any site effects, site was included as a random effect in applicable statistical models (see statistical analysis section for details).

In the lab, oysters were cleaned and prepared over eight days while maintained in 50-L flow-through tanks. To measure the carbonate chemistry of the EPF, a 2mm hole was drilled into the right valve approximately two centimeters from the hinge to expose the EPF cavity, without damaging the underlying mantle tissue. The drilled hole was gently rinsed with filtered seawater and patted dry. The barbed end of a nylon luer-lock coupling (McMaster-Carr 51525K123) was trimmed to the same length as the shell thickness and inserted into the hole and attached using marine-safe cyanoacrylate (Starbond EM-2000 CA USA). The coupling was then plugged with a nylon cap socket (McMaster-Carr 51525K315). Oysters were allowed to recover for 4 days, after which they were transferred to the experimental tank array (described below) for acclimation and exposure. During the transfer, oysters were assigned one of three *p*CO_2_ treatments: ‘control’ (ca. 500 uatm) – corresponding to near-present-day; ‘moderate OA’ (ca. 1000 uatm) – corresponding end-of-century predictions (Intergovernmental Panel on Climate Change, 2007); and ‘high OA’ (ca. 2800 uatm) – corresponding to seawater conditions that are undersaturated with respect to the oyster shell calcite (Ω_calcite_ < 1). Importantly, high OA events are already regularly experienced in the coastal marine systems from which these oysters were collected, owing to daily tidal cycles, input of meteoric waters, and seasonal reoxidation of organic matter (Figure S1.1). This trend is underappreciated in many coastal systems, where local processes are increasing the acidity of the seawater relative to the global average (Reum et al., 2014). The high OA treatment was also utilized to increase the inferential and statistical power of the study design (Whitlock and Schluter, 2014).

### Experimental design

The experiment was conducted at Northeastern University’s Marine Science Center, using a flow-through seawater system that draws water from the Broad Sound, adjacent to Nahant, Massachusetts (42.416884, −70.907564). Each *p*CO_2_ treatment was replicated over two blocks, each containing three 42-L acrylic aquaria (total of 6 aquaria per treatment). Tanks within a block were connected via the recirculation and filtration system, while gas mixture, temperature, and incoming filtered seawater were regulated independently for each tank. A blocked randomization design was used to ensure an equal number of oysters from each location were distributed across all treatments and that there were no significant differences in starting oyster size across treatment (mean±SD shell length: 9.70±2.4cm). Twelve oysters were placed into each aquarium, four from each location, resulting in 24 oysters per treatment per location (total of 72 oysters per treatment). Oysters were acclimated for 23 days at present-day pCO_2_ conditions at a temperature of 17 C. Following acclimation, the medium- and high-OA treatments were ramped up to target *p*CO_2_ over a 12-hour period. Treatment levels were then maintained for 80 days.

Target *p*CO_2_ gases were formulated by mixing compressed CO_2_ with compressed CO_2_-free air (low-OA treatment) or with compressed air (moderate- and high-OA treatments) using solenoid-valve-controlled mass flow controllers (Aalborg mass flow controllers, Model GFC17, precision = 0.1mL/min) at flow-rates proportional to the target pCO_2_ conditions. The CO_2_-free air used in the low-OA treatment was generated by scrubbing CO_2_ from compressed air with a Parker Balston FT-IR Purge Gas Generator. The mixed gases were then bubbled into the experimental seawater treatments with 45-cm flexible microporous tubes at rates sufficient for the pCO_2_ of aquaria seawater to equilibrate with the pCO_2_ of the mixed gases. Filtered seawater was introduced to each aquarium at a flow rate of 150 mL min^−1^. Temperature of all experimental tanks was maintained at 17°C with 1/4HP chillers (Aqua Euro USA, precision = 0.1°C). Seawater temperature of all aquaria was slowly increased to 18.5°C (Figure S1.2) on days 35-37 of the experiment in an effort to stimulate gonad development for a companion experiment. It should be noted that this is a small temperature shift relative to what these oysters experience seasonally in their native habitats (Figure S1.1). Oysters were fed 1% Shellfish Diet 1800^®^ twice daily following best practices outlined in (Helm and Bourne, 2004).

Oyster EPF and tissue was sampled at six discrete time-points that decreased in frequency throughout the duration of the experiment (time post 12 hour ramp up period: 24 hours, 48 hours, 9 days, 22 days, 50 days, and 80 days). This sampling schedule was designed to capture rapid physiological changes that occurred during the onset of exposure and to examine long-term stability of physiological and molecular responses.

### Measurement of the seawater carbonate system

Temperature, pH, and salinity of all tanks were measured three times per week (M, W, F) for the duration of the experiment. Seawater pH was measured with an Accumet solid state pH electrode (precision = 1mV), salinity was measured using a YSI 3200 conductivity probe (precision = 0.1 ppt), and temperature was measured using a NIST-standardized glass thermometer (precision = 0.1 °C). Seawater samples were collected every two weeks from each tank for analysis of dissolved inorganic carbon (DIC) and total alkalinity (TA) on a VINDTA 3C coupled alkalinity gram titration and coulometric DIC analyzer system. In brief, samples were collected in 250 ml borosilicate glass bottles and immediately poisoned with 100 ul saturated HgCl_2_ solution, then refrigerated until analyzed. DIC, TA, salinity, and temperature were used to calculate calcite saturation state, pH, CO_3_^2-^, HCO_3_^-^, aqueous CO_2_, and pCO_2_ of each sample using CO_2_SYS version 2.1 (Pierrot et al., 2011), using the seawater pH scale with K1 and K_2_ values from (Roy et al., 1993), a KHSO_4_ value from (Dickson, 1990), and a [B]_T_ value from (Lee et al., 2010).

### Extrapallial fluid chemistry

Oyster pH_EPF_ was measured by removing each oyster from their tank, inserting a 5 mL syringe with a flexible 18-gauge polyproplyene tip through the luer-lock port into the oyster’s extrapallial cavity and extracting approximately 0.5-2 mL of fluid. Care was taken to avoid puncturing the mantle tissue and inadvertently sampling either the hemolymph or stomach fluid. The pH_EPF_ was measured immediately after extraction with an Orion 91’10DJWP Double Junction micro-pH probe standardized with pH 7.01 and 10.01 NBS buffers.

### Calcification rate

Net calcification rate was calculated for oysters surviving to either 50 or 80 days (*n* = 35) by buoyantly weighing oysters prior to exposure (BW_1_) and on day 33 or 34 of the exposure (BW_2_) following the methods of Ries at al. (2009). Buoyant weight was measured in a 27.65 liter tank (48 cm long, 24 cm wide and 24 cm deep) filled with seawater from the flow-through system and was maintained at treatment temperature by an Aqua Euro USA Model MC-1/4HP aquarium chiller. Buoyant weight was measured by completely submerging the oyster on a flat platform suspended from a bottom-loading scale (Cole Parmer Symmetry S-PT 413E, precision = 0.001 g). Care was taken to ensure no bubbles were trapped inside oyster shells. Oysters were weighed three times in the weighing basket, removing the oyster from the basket between each measurement. If replicate measurements varied by more than 0.01 g, oysters were re-weighed. A standard of known weight was weighed every 20 oysters to ensure that no drift was occurring in the scale.

To establish a relationship between buoyant weight and dry weight for the purpose of estimating net calcification rate, shells of oysters sampled for tissue within four days of a buoyant weight measurement were soaked in 10% ethanol to remove salts, dried, and weighed. The dry weight was then regressed against the buoyant weight measurement to establish an empirical dry-buoyant weight relationship (Figure S2.1):

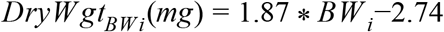

This empirical relationship was then used to calculate dry shell weight at each buoyant weight time point via linear regression.

Calculated dry weights were then used to calculate daily calcification rate:

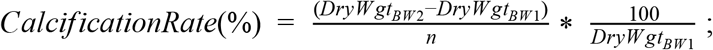

where dry weight (DryWgt_BWi_) was calculated for each individual using the buoyant weight pre-exposure (BW1) and 33-34 days into the exposure (BW2) and *n* was the number of days between the two measurements. Lastly, the average daily change in dry weight was divided by initial dry weight to standardize calcification rate for allometric effects, and multiplied by 100 to convert that fraction into a percent.

### Tissue collection

Immediately following extraction of EPF at each time point, oysters were shucked and the edge of the mantle tissue was sampled. Patterns observed in the response of pH_EPF_ over time informed the decision to focus genetic analyses on mantle edge tissue extracted on days 9 and 80 of the exposure (6 from low-OA and high-OA treatments at both time points, n = 24). For consistency and to minimize potential effects on mantle tissue arising from drilling of the oysters’ right valves when installing the EPF extraction ports, mantle tissue was only collected from the left valve of the oysters (i.e., the side opposite of the EPF extraction port). Tissue was flash frozen in liquid nitrogen before being transferred to a −80 °C freezer for storage prior to DNA/RNA extraction.

### DNA methylation library preparation and quantification

DNA was isolated from 24 oyster mantle edge tissue samples using the E.Z.N.A. Mollusc Kit (Omega) pursuant to the manufacturer’s instructions. Isolated DNA was quantified (Qubit dsDNA BR Kit; Invitrogen, CA, USA). DNA samples were sonicated for 10 minutes at 4 °C, at 30 second intervals at 25% intensity with a ME220 Focused-ultrasonicator (Covaris, MA, USA). Shearing size (350bp) was verified using a 2200 TapeStation System (Agilent Technologies, Santa Clara, CA, USA). Samples were enriched for methylated DNA using MethylMiner kit (Invitrogen, CA, USA). Enriched samples were then sent to Zymo Research (Zymo, CA, USA) for bisulfite-conversion, library preparation, and sequencing. Libraries were sequenced on an Illumina HiSeq1500 platform with paired-end 100bp sequencing.

Raw sequences were trimmed to remove adapters and low quality sequences by removing 10 bp from both 5’ and 3’ ends of each fragment using the command *trim_galore* in the program TrimGalore! (Martin, 2011) with the *--clip_r1, --clip_r2, --three_prime_clip_R1*, and *--three_prime_clip_R2* flags set to 10, and the remaining flags left as the defaults. Quality of sequences was assessed with FastQC (Leggett et al., 2013) within the *trim_galore* command using the *--fastqc_args* flag. Next, a bisulfite converted version of the *C. virginica* genome (NCBI Accession GCA_002022765.4) was prepared with the *bismark_genome_preparation* command in Bismark (Krueger and Andrews, 2011) and the --bowtie2 flag. The trimmed sequences were then aligned to the prepared reference using the *bismark* command with *--non-directionality* specified and the alignment score set to *-score_min L,0,-0.8*. Duplicates were removed from the data using the command *deduplicate_bismark*. All cytosines in the genome with coverage were extracted from the aligned deduplicated data for each individual using the command *bismark_methylation_extractor* with default settings. Next, a coverage report for all CpG associated cytosines within the genome was generated for each individual with the command *coverage2cytosine* using the output from the previous step. This report contains all cytosine loci (even those with no coverage) located within a CpG motif and includes separate columns for methylated vs. unmethylated coverage. Files were then processed using a standard methylKit pipeline, which included normalization of methylation counts among samples, destranding CpGs loci, and filtering loci that did not have at least 5x coverage for each sample (Akalin et al., 2012). Here, destranding involves combining the cytosine calls within a single CpG from either strand (i.e., top or bottom) and the proportion methylation was determined by dividing the number of methylated cytosine calls by the total coverage (regardless of methylation status). Finally, CpG loci were annotated by feature (e.g., introns, exons, and intergenic regions) and genomic coordinate using feature tracks created based on methods outlined in (Venkataraman et al., 2020) using the gene annotation file available on NCBI (Accession: GCA_002022765.4). Hereafter, ‘gene body’ is used to reference the transcriptional region of the gene, which includes exons, introns, and untranslated regions, but not promoters.

### RNA library preparation and quantification

RNA was extracted from the same 24 samples used for DNA methylation using TRI Reagent Protocol (Applied Biosystems) following manufacturer’s instructions. A cDNA library was constructed for each sample using a TruSeq Stranded mRNA library Prep kit (Illumina, Cat# RS-122-2101) following manufacturer’s instructions. Library fragment size was determined using a BioAnalyzer (RNA 6000 Nano Kit; Agilent, CA, USA) and concentration was quantified on a Qubit (Qubit High Sensitivity RNA Kit; Invitrogen, CA, USA). Samples were sent to Genewiz (NJ, USA) for library preparation and sequencing. Libraries were sequenced on two lanes (12 individuals/lane) of an Illumina HiSeq 3000 platform with paired-end 200bp sequencing.

RNA reads were first trimmed to remove adapters and for quality control using Trimmomatic (Bolger et al., 2014), implemented in the dDocent pipeline (v.2.2.20; Puritz et al., 2014) and following recommendations by Puritz et al. (2014). This was done using the *dDocent* command with default settings, which included removing adapters and performing a quality control step that trimmed the leading and trailing bases with phred quality scores < 20, with additional trimming of 5bp windows with mean phred quality scores < 10 (see github repository for complete description). Reads were then aligned to the *C. virginica* genome (Accession: GCA_002022765.4) using a two-step mapping approach with the program STAR (v.2.7.0; Dobin et al., 2013). In the first step, a preliminary alignment to the reference genome was performed for each sample to identify novel splice junctions. In the second step, reads were realigned, but with all the splice junctions discovered from step one included along with the reference genome. Both steps were executed with the command *STAR* using default settings, with the exception that the number of alignments retained was adjusted by setting both the *--outFilterMatchNminOverLread* and *--outFilterScoreMinOverLread* flags to 0.17, and increasing the number of discoverable sequence junctions by setting the flag *--limitSjdbInsertNsj* to 1500000. In the second step, the *--sjdbFileChrStartEnd* flag was also included to specify the inclusion of splice junctions identified in step one. This two-step alignment process has been shown to improve mapping quality (Dobin et al., 2013).

Expression levels for each gene were quantified for each sample with the program RSEM (Li and Dewey, 2011) using the *rsem-calculate-expression* command with *--star* flag, default settings, the bam files generated from the mapping step with STAR, and the same *C. virginica* gene annotations used during mapping (Accession: GCA_002022765.4). Importantly, RSEM takes a probabilistic approach to transcript quantification, which allows for the retention and fractional assignment of multi-mapping reads. Single sample gene count estimates were combined into count matrices (number of samples x number of genes) using a custom shell script (available on the supplemental github repository), then filtered to remove genes that did not contain at least 1 count per million transcripts in at least five individuals within one of the treatment-time experimental levels.

### Statistical Analyses

All statistical analysis was performed in R (v3.6.0, R Core Team 2019), using the graphical user interface RStudio (v.1.2.1335, RStudio Team 2018). See Data Accessibility for R scripts.

### Extrapallial fluid pH (pH_EPF_) and calcification

A linear mixed model was used to determine the effect of the explanatory variables of treatment and time on the response variable of pH_EPF_. The full model included the explanatory variables of treatment and time and their interaction as categorical fixed effects, and tank (nested within shelf) and oyster collection site as random effects. The model was performed in R using the *lme4* package (Bates et al., 2015). A step-down strategy with likelihood ratio tests (based on the degrees of freedom estimated using the Satterthwaite method) implemented in *lmerTest*, following the principle of marginality (Kuznetsova et al., 2017), was used to select the most parsimonious linear mixed effects model. This model selection approach evaluates two different interpretations of pH_EPF_: (1) the measured value of pH_EPF_ and (2) pH_EPF_ relative to treatment seawater pH (ΔpH = pH_EPF_ - pH_seawater_), which may be a better indicator of active pH_EPF_ regulation. To evaluate significant differences between pH_EPF_ and ΔpH under the moderate and high OA treatments for each time point, a series of post hoc comparisons with a Tukey correction were performed using the *multcomp* package (Hothorn et al., 2008). In addition, the change in ΔpH over time was examined by performing a series of single sample t-tests to determine whether each treatment at each time point was significantly different from seawater pH. We corrected for testing multiple hypotheses using a Benjamini-Hochberg correction (Benjamini and Hochberg, 1995).

Linear mixed effect models were used to examine the effect of pCO_2_ on long term EPF response, both measured pH_EPF_ and ΔpH. The full models included EPF response (pH_EPF_ or ΔpH) as the response variable, treatment and time and their interaction as categorical fixed effects and tank (nested within shelf) and oyster collection site as random effects. We included samples from our two long term time points (day 50 and 80) in the model (n = 35). Model selection and post hoc comparisons were performed using the approach described above.

Another set of linear mixed effect models were used to examine the effect of either treatment or pH_EPF_ on calcification rates. In the full models, tank (nested in shelf) and collection site were included as random effects. Calcification rate was based on buoyant weights measured at a single time (day 33) for all oysters (n = 35), so time could not be included as a fixed effect. Treatment was handled as a continuous variable and was calculated as the average pCO_2_ for each tank across the duration of the exposure prior to sampling. Model selection was performed using the approach described above. Regression was used to examine the effect of either treatment or pH_EPF_ and calcification rate.

### Genome-wide analysis

#### Global median methylation analysis

A linear model including treatment and time and feature (i.e., exon, intron, and intergenic region) as explanatory factors was used to examine global patterns in median methylation among treatments and across time. Only CpG loci with at least 5x coverage for all samples were included.

#### Principal components analysis and discriminant analysis of principal components

A principal components analysis (PCA) was used to visualize differences amongst treatments and time points in both global gene expression and gene body DNA methylation patterns. The gene expression PCA was based on the post-filtered expression level of genes (in columns) for each individual (in rows). Gene expression level was calculated as counts per million using the *cpm* function from the R package edgeR (Robinson et al., 2010) and log2 transformed (i.e., log2-cpm). The DNA methylation PCA was based on all CpG loci located within gene bodies with at least 5x total coverage (in columns) for each individual (in rows).

We used a PERMANOVA to test the null hypothesis of no effect of treatment, time, or their interaction on global gene expression and DNA methylation patterns. The PERMANOVA was based on the Manhattan distance using the *adonis* function in the R package *vegan* (v2.5-5; Dixon, 2003)

To further investigate the specific effect of OA on genome-wide variation in both differential gene expression and DNA methylation, a discriminant analysis of principal components (DAPC) was performed with the R package adegenet (Jombart et al., 2010), using the same transcriptomic and methylomic datasets described above for the PCA. In brief, this method defines a discriminant function that maximally differentiates between two or more categories based on multidimensional sample data. Here, a DAPC was used to generate a discriminant function that maximized differences between treatments in day 9 samples using the function *dapc*, then we predicted where samples from day 80 should fall along the discriminant function using the *predict* function to determine if genome-wide variation due to OA was maintained through time.

### Differential molecular response analysis

#### Identification of differentially methylated loci

Differentially methylated loci (DML) were identified using the R package methylKit (Akalin et al., 2012). Only CpGs with coverage ≥ 5 for all samples were considered. Differential methylation was performed using a logistic regression approach implemented in methylKit with the functions *calculateDiffMeth* with the overdispersion argument set to “MN” and the default ‘slim’ method to correct p-values and the function *getMethylDiff* with a differential methylation threshold set to 50% and a q-value threshold set to 0.01.

#### Identification of differentially expressed genes

Differential gene expression amongst treatments was evaluated using a generalized linear model approach implemented in the R package *limma* (Ritchie et al., 2015) using treatment, time, and their interaction as fixed effects. Expression data was first TMM normalized using the *calcNormFactors* function and transformed into log2 counts per million (log2-cpm) using the *voomWithQualityWeights* function (Smyth et al., 2005). Finally, the *geneDuplication* function was used to account for tank as a potential experimental block effect (Oshlack et al., 2007). Site was not considered in this analysis given that it did not have a significant effect on either the phenotypic or genome-wide responses. Genes with FDR ≤ 0.05 and absolute value of log_2_ fold ≥2 were considered differentially expressed.

### Gene co-expression network analysis

A weigheted gene co-expression network analysis was performed to identify genes that exhibit similar expression patterns among individual oysters using the R package WGCNA (Langfelder and Horvath, 2008). First, a gene dissimilarity matrix was generated based on the log2-cpm gene expression data using first the *adjacency* function followed by the *TOMsimilarity* function in WGCNA. This step estimates the level of dissimilarity between each gene by considering expression across all individuals. Next, genes were hierarchically clustered based on dissimilarity using the function *hclust* and the ‘Ward.D2’ method for clustering (Murtagh and Legendre, 2014). Modules were determined using the *cutreeDynamic* function with a minimum gene membership threshold of 30. An eigenvalue for module expression (i.e., the first principle component value for each individual) was calculated for each module using *moduleEigengenes*. Lastly, linear regression was used to determine the association between the expression of each module (i.e., the eigenvalue of gene expression) and either mean gene methylation (calculated as the mean methylation of all CpGs among all genes within a module) or EPF response (i.e., ΔpH).

### Functional enrichment analysis

A functional enrichment test was conducted with GO-MWU, a rank-based gene enrichment method developed by (Wright et al., 2015), to identify gene ontology (GO) categories enriched among genes that are differentially regulated or methylated between treatments at each time point. We performed this analysis separately for each time point using the log_2_-fold change in gene expression and the difference in mean methylation among treatments. Mean methylation was calculated as the mean among all CpG loci within a gene across all individuals within a particular treatment and time point. Only genes with at least 5 CpG loci were considered for the analysis to ensure mean methylation estimates were based on genes where we had at least moderate CpGs coverage. Importantly, GO-MWU can handle a variety of differentiation metrics (e.g., log_2_-fold change in expression) and considers all genes, not just those that are significantly differentially expressed or methylated. This enables detection of GO categories enriched with responsive genes even when there is limited evidence of individually differentially expressed or methylated genes. GO-MWU scripts and the gene ontology database were downloaded from the GO-MWU github repository (https://github.com/z0on/GO_MWU).

Inputs for the GO-MWU analysis include two gene list tables created using the Genebanks IDs from the *C. virginica* genome available on NCBI (Accession: GCA_002022765.4) along with a measure of difference (i.e., log_2_-fold change or methylation difference) and a table of GO terms containing a list of all available Genebank IDs and their associated gene ontology (GO) terms. Details on how the latter file was created are outlined by Johnson et al (2019) and can be found on the associated Github repository (See data availability section). The GO-MWU analysis was performed for both gene expression and methylation tables separately using the *goStats* function in R with default settings and using the gene ontology database provided by the GO-MWU repository. The analysis first clusters highly similar GO categories by combining categories that shared at least 75% of the same genes. After clustering, a Mann-Whitney U test was performed to identify GO categories that were enriched with either up-regulated or down-regulated (or hyper- or hypo-methylated) genes. This analysis was run separately for GO categories associated with molecular function, biological process, and cellular components. A 10% FDR correction (GO-MWU default) was used to adjust for multiple comparisons.

### Gene-level characterization of gene expression and DNA methylation

A PCA-based approach (described in Gavery and Roberts, 2013) was used to examine the relationship between gene expression, DNA methylation, and gene attributes (e.g., gene length and number of exons). The following variables were used in the input matrix for the PCA: mean gene expression over all treatments (mean log2-cpm values); the coefficient of variation (CV) in gene expression among treatment means; mean DNA methylation level over all treatments; the DNA methylation CV among treatment means, gene length, exon number per gene; and the number of CpGs per gene (script for generating each variable within the matrix are available on the github repository). Gene attributes were normalized by log transformation.

## Results

### Experimental seawater chemistry

After a 23 day acclimation period (pCO_2_ 547.3 ±20.2 ppm), tanks were incrementally adjusted to the target pCO_2_ of the three experimental treatments (mean±SEM, control: pCO_2_ 579.1 ±16.5 ppm, moderate OA: pCO_2_ 1050.4±47.5 ppm, high OA: pCO_2_ 2728.6± 128.0 ppm, Table S1.1).

### Extrapallial fluid chemistry and calcification response

The best model to explain pH_EPF_ included both fixed effects (treatment and time), their interaction, and the random effect of tank nested within shelf. There was a significant effect of treatment (F_2,14.85_ = 4.936; *P* = 0.0227) and the interaction between time and treatment (F_10,73.985_ = 2.38; *P* = 0.017) for pH_EPF_ evaluated across all 6 time points (*n* = 107). Post-hoc tests revealed a significant decrease in pH_EPF_ within the high OA treatment compared to the control condition at the onset of the exposure (24-48 hours), no change on days 9-22, and a significant decrease between the final two time points (day 50 and 80; Figure 1A).

**Fig. 1.**
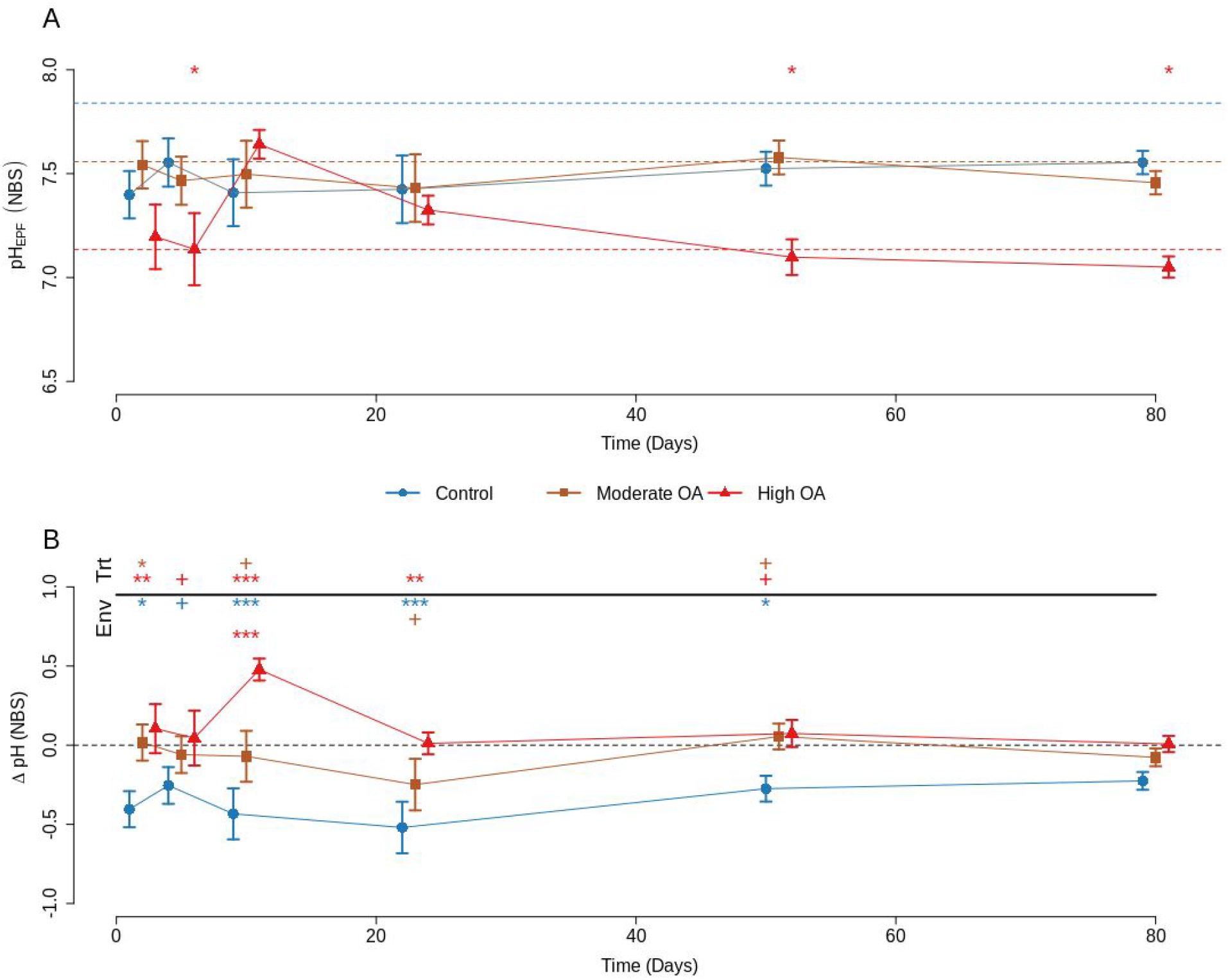
EPF pH over the 80 day exposure. **(A)** pH_EPF_ and **(B)** ΔpH (pH_EPF_ - pH_seawater_) across time with standard error bars. Dotted lines represent the pHseawater through time. In **(A)** colored dashed lines represent each treatment level averaged over the duration of the exposure and symbols at the top of the graph indicate if the pH_EPF_ in one of the OA treatments (‘moderate OA’, 1000 uatm; ‘high OA’, 2800 μatm) was significantly different from the pH_EPF_ in the control condition (‘control’, 580 μatm) for each time point (e.g., H_0_: control pH_EPF_ = high OA pH_EPF_; top row: moderate OA, bottom row: high OA). In **(B)** the black dashed line represents the local 2 week average for each sample point which has been standardized to zero. Symbols at the top of the graph indicate significant post hoc tests, comparing the OA treatments to the control (i.e., ‘Trt’) or all treatments to the pHseawater (i.e., ‘Env’). In the ‘Trt’ comparisons, symbols indicate time points where either OA treatment was significantly different from the control (e.g., H_0_: control ΔpH= high OA ΔpH; top row: moderate OA, bottom row: high OA). In the ‘Env’ comparisons, symbols indicate time points where any treatment was significantly different from 0 (top row: Control, middle row: moderate OA, bottom row: high OA). Asterisks (*** = *P* < 0.001, * = *P* < 0.05) and pluses (+ = *P* <0.1) indicate degree of significance. Treatment points within time points were staggered along the x axis to improve visualization.

The best model to explain ΔpH (pH_EPF_ - pH_seawater_) included both fixed effects (treatment and time), their interaction, and the random effect of tank nested within shelf. Treatment had a significant effect (F_2,14.85_ = 13.412; *P* = <0.001), while the effect of time (F_5,73.993_ = 2.097; *P* = 0.075) and the interaction of treatment and time (F_10,73.985_ = 1.87; *P* = 0.063) were not significant (Figure 1B). Post-hoc comparisons between the control treatment and each OA treatment at each time point revealed a significant increase in ΔpH within the high OA and to some extent the moderate OA treatments compared to the control condition at the onset of the exposure (24-22 days). By days 50 and 80, ΔpH was no longer statistically different amongst treatments, despite substantially higher mean ΔpH in both OA treatments relative to the control (Figure 1B, ‘Trt’). One-tailed t-tests showed that ΔpH of oysters in the control treatment was significantly lower than 0 (i.e., the EPF was more acidic than the pH_seawater_) at almost all time points of the experiment (Figure 1B, ‘Env’), indicating a strong tendency for oysters to maintain a more acidic EPF fluid relative to the environment in control conditions. In contrast, pH_EPF_ of oysters in the elevated pCO_2_ treatments did not significantly differ from seawater pH at most time points, with the exception of day 9 in the high OA treatment when ΔpH was significantly higher than 0 (Figure 1B).

The best model to explain pH_EPF_ and ΔpH over the long-term (i.e., conditions on days 50 and 80; n = 35), included treatment as a fixed effect. Treatment was a significant predictor of pH_EPF_ (F_2,32_ = 23.85, *P* < 0.0001) and ΔpH (F_2,32_= 8.04, *P* = 0.0015; Figure 2A-B). The post hoc tests revealed a significant decrease in pH_EPF_ in the high OA treatment relative to both the control and moderate OA treatments, while post hoc tests showed an increase in ΔpH in the two OA treatments relative to the control.

**Fig. 2.**
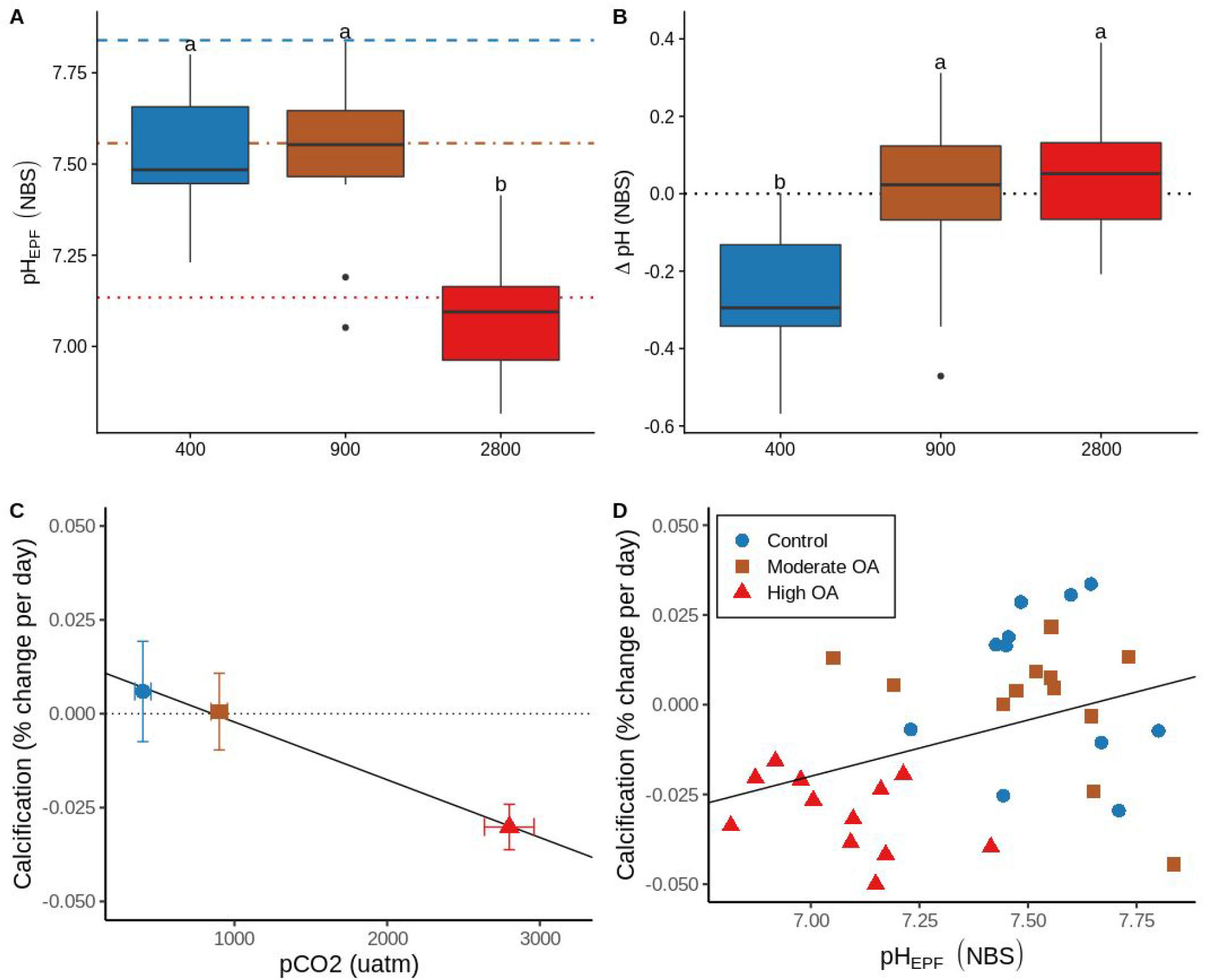
Long term trends in EPF pH and calcification. Boxplots of long term (**A**) pH_EPF_ by treatment, horizontal lines indicate mean seawater pH for the three treatments (dashed: control, dot-dashed: moderate OA, dotted: high OA), and (**B)** ΔpH (pH_EPF_ - pH_seawater_) by treatment, dotted line at zero represents the mean adjusted pH_seawater_. (**C**) Percent daily calcification rate by measured treatment pCO_2_ (*slope* = −1.940e-07, *P* < 0.001, R^2^ = 0.465), dotted line indicates calcification rate of zero percent and solid black line represents fitted regression. (**D**) Percent daily calcification rate by pH_EPF_ (*slope* = 0.031, *P* = 0.026, R^2^ = 0.114), solid black line represents fitted regression. EPF and calcification measurements based on individuals from either day 50 or 80 time points (n = 35). Letters on on first two panels (**A** and **B**) represent levels of significance based on post hoc testing, while bars on (**C**) represent standard errors.

The best model to explain calcification rate response to environmental pCO_2_ included treatment as the only fixed effect. Calcification rate (*n* = 35) significantly decreased with increasing OA (% change per ppm_CO2_, R^2^ = 0.465, *P* = <0.001). Furthermore, this pattern was driven largely by the decrease in calcification rate in the high OA treatment (Figure 2C), where net calcification rate decreased below 0. This indicates that shell dissolution was occurring in the high OA treatment. Lastly, a linear regression model that evaluated the correlation between calcification response and pH_EPF_ (with pH_EPF_ as an explanatory variable) found a significant positive relationship between pH_EPF_ and calcification rate (R^2^ = 0.114, *P* = 0.026, Figure 2D).

### DNA methylation responses to OA

Approximately 1.4 billion paired-end 80bp reads of MBD-enriched, bisulfite treated DNA were obtained across the same 24 samples used for RNA sequencing (60.0±8.4 million reads per individual, NCBI BioProject ID: PRJNA594029, Table S3.1). A total of 622.4 million quality filtered reads mapped to the *C. virginica* genome (mapped to 12,765,452 CpGs of a total of 14,458,703 CpGs in the genome, 88.3% of all CpGs in the genome). A single individual within the control treatment on day 9 was removed from downstream analysis due to poor sequencing depth and a high rate of gene duplication error after mapping. After filtering, we retained 403,976 CpGs with at least 5x coverage for each of the remaining 23 samples (Table S3.2). Of the CpGs with at least 5x coverage, 380,168 (94.1%) were located within gene bodies, with 242,102 CpGs located in exons among 27,932 genes (Figure 3A “CpGs_5_”). CpGs with a minimum of 5x coverage in each sample were used for the downstream analyses.

**Figure 3.**
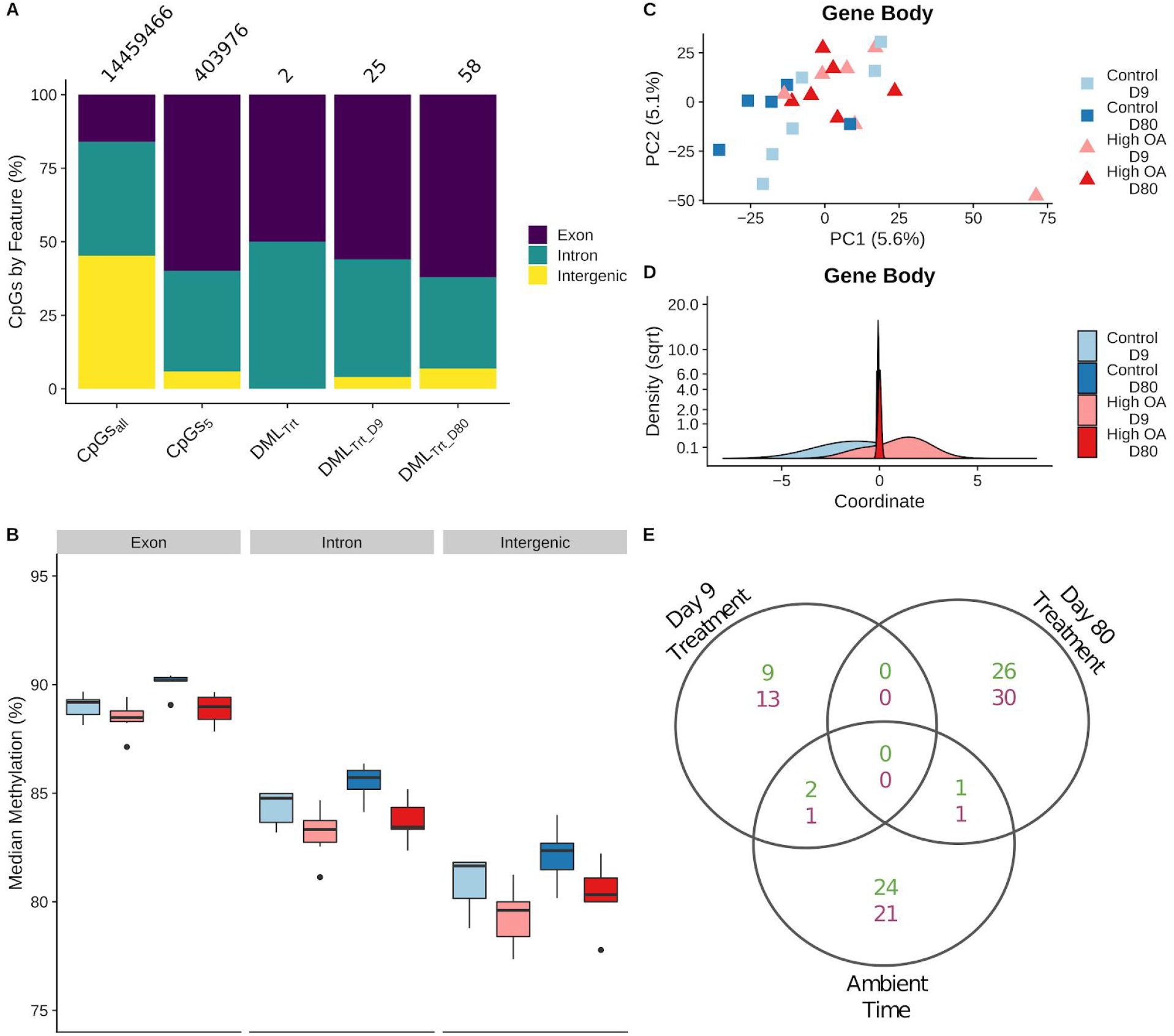
DNA methylation responses to OA. **(A)** Proportion of CpGs by feature across data subsets. CpGs_all_: all CpGs in genome; CpGs_5_: CpGs with at least 5X coverage for each individual; DML_Trt_: differentially methylated loci (DMLs) between control and OA treatments across both time points; DML_Trt_09_ and DML_Trt_80_: DMLs between treatments on day 9 and 80, respectively. Numbers above the bars represent the total number of CpGs for each group. **(B)** Boxplot of median global methylation for each sample by genome feature (*P_Tr_*_t_ < 0.0001, *P_Time_* = 0.001, *P_Feature_* < 0.0001). **(C)** Plot of the first two principal components from a principal components analysis (*P_Trt_* = 0.027, *P_Time_* = 0.235, *P_TimexTrt_* = 0.364), and (**D**) density plot of the discriminant values from a DAPC based on gene body methylation (exons and introns). Discriminant values were determined by a function that maximally discriminates between treatment using samples from day 9, then the discriminant value for each sample from day 80 was predicted using the same discriminant function. **(E)** Venn diagram of DMLs among treatments for each day (Day 9 Treatment and Day 80 Treatment) and among time points in the control treatment (Control Time). Overlapping regions indicate DMLs shared among comparisons. Hypermethylated DMLs in green (top number), hypomethylated DMLs in purple (bottom number).

An ANOVA examining global median methylation (median percent methylation for all CpGs within a feature for an individual) found significant effects of treatment, time, and feature, but not their interactions (Figure 3B, *P_Trt_* < 0.0001, *P_Time_* = 0.001, *P_Feature_* < 0.0001). Genome-wide DNA methylation was the most different among features, with exons exhibiting the highest levels of methylation. We also found that OA led to significant hypomethylation, while time led to significant hypermethylation, although these only led to a 1-5% shift in methylation when averaged over all loci in the genome.

A PERMANOVA considering genome-wide gene body methylation for all individuals revealed significant global methylation differences due to treatment (*Adonis P_Trt_* = 0.027), but not by time (*Adonis P_Time_* = 0.235) or their interaction (*Adonis P_TimexTrt_* = 0.364; Figure 3C). Notably, a DAPC using the same gene body CpGs showed only moderate separation in global DNA methylation among treatments at day 9, and the variation that maximally differentiated the treatments at day 9 was not maintained by day 80 (Figure 3D), suggesting that CpG loci involved in differentiating between treatments at day 9 may not be the same as those on day 80.

A logistic regression approach implemented in methylKit was used to identify DML. The initial model included treatment as the main effect and time as a covariate (DML_Trt_). Next, three follow up models were used to examine the effect of treatment for each time point in separate comparisons (day 9: DML_Trt_D09_; day 80: DML_Trt_D80_) and the effect of time using samples from the control treatment. Only 2 CpGs were found to be differentially methylated by treatment, both located in gene bodies (Figure 3A, “DML_Trt_”). Follow up comparisons found additional DML by treatment on either day 9 (25) or day 80 (58) (Figure 3A, “DML_Trt_D09_” and “DML_Trt_D80_”), with no overlap among the two groups of DML (Figure 3E), further illustrating that the loci being differentially methylated in response to OA shifted through time. A shift in methylation at specific loci was also observed through time in the control treatment, although there was little overlap between these DML and those responding to OA (Figure 3E). Most of the DML by treatment identified on either day were located within gene bodies (day 9: 96%, day 80: 93.1%), with only a few genes containing multiple DML (day 9: 9.1%, day 80: 5.8%). We saw no overlap of specific DML by treatment among days. We found multiple DML on both days in genes associated with the THO complex, a nuclear structure comprised of multiple proteins involved in transcription elongation and mRNA maturation, including hypermethylated DML in THO complex subunit 1 on day 9 and hypomethylated DML in THO complex subunit 2 on day 80 (Table S3.3). On day 80, we also found DML in putative cadherin, protein ubiquitination, and death effector domain-containing genes. Two GO categories were enriched in hypomethylated CpGs, the cellular component ‘amidotransferase complex’ on day 9 and the biological process ‘biosynthetic process’ on day 80 (Table 1). No categories were shared among time points.

**Table 1.** Gene ontology (GO) categories enriched in differentially responsive genes by treatment, summarized by molecular function (MF), biological process (BP), and cellular component (CC). Up- and down-regulation refers to the change in gene expression (GE) in the high OA treatment relative to the control treatment (i.e., the GO categories enriched in up-regulated genes had higher gene expression under high OA compared to the control treatment). Similarly, hypomethylation indicates the GO category was enriched with CpG loci (DNAm) that were less methylated under high OA relative to the control treatment. Names in bold represent GO categories enriched at both time points. The full category name for “oxidoreductase activity *” is “oxidoreductase activity, acting on paired donors, with incorporation or reduction of molecular oxygen”.

| Gene Expression |  |  |  |  |  |
| --- | --- | --- | --- | --- | --- |
| Summary Category | Day | Direction in OA Treatment | Name | Delta Rank | P (adj) |
| BP | 9 | Down regulated | signal transduction | -192 | 0.0058 |
| BP | 9 | Up regulated | biosynthetic process | 235 | 0.0058 |
| BP | 9 | Up regulated | **cellular nitrogen compound metabolic process** | 164 | 0.0315 |
| BP | 9 | Down regulated | G protein-coupled receptor signaling pathway | -227 | 0.0315 |
| BP | 9 | Up regulated | **DNA metabolic process** | 336 | 0.0334 |
| BP | 80 | Down regulated | macromolecule modification | -374 | <0.0001 |
| BP | 80 | Down regulated | dephosphorylation | -523 | <0.0001 |
| BP | 80 | Down regulated | phosphorus metabolic process | -288 | <0.0001 |
| BP | 80 | Up regulated | **DNA metabolic process** | 421 | 0.0008 |
| BP | 80 | Down regulated | protein metabolic process | -200 | 0.0012 |
| BP | 80 | Up regulated | **cellular nitrogen compound metabolic process** | 178 | 0.0054 |
| BP | 80 | Up regulated | DNA recombination | 532 | 0.0211 |
| BP | 80 | Up regulated | glutamine family amino acid metabolic process | 535 | 0.0281 |
| BP | 80 | Up regulated | glutamate biosynthetic process | 1676 | 0.0443 |
| BP | 80 | Up regulated | dicarboxylic acid biosynthetic process | 1237 | 0.0443 |
| BP | 80 | Up regulated | macromolecule biosynthetic process | 263 | 0.0474 |
| CC | 9 | Down regulated | **transcription factor complex** | -147 | 0.0170 |
| CC | 9 | Up regulated | ribosome | 169 | 0.0431 |
| CC | 9 | Up regulated | intracellular non-membrane-bounded organelle | 139 | 0.0454 |
| CC | 80 | Down regulated | **transcription factor complex** | -159 | 0.0045 |
| CC | 80 | Down regulated | protein-containing complex | -105 | 0.0154 |
| MF | 80 | Up regulated | DNA binding | 352 | 0.0245 |
| MF | 80 | Up regulated | RNA-DNA hybrid ribonuclease activity | 1320 | 0.0245 |
| MF | 80 | Up regulated | oxidoreductase activity | 219 | 0.0245 |
| MF | 80 | Up regulated | oxidoreductase activity * | 321 | 0.0245 |
| MF | 80 | Up regulated | double-stranded DNA binding | 905 | 0.0487 |
| DNA Methylation |  |  |  |  |  |
| BP | 80 | Hypomethylated | cellular modified amino acid biosynthetic process | -4437 | 0.0333 |
| CC | 9 | Hypomethylated | glutamyl-tRNA(Gln) amidotransferase complex | -8528 | 0.0179 |

### Gene expression responses to OA

RNA sequencing yielded a total of 955 million paired-end reads (40±2.7 million reads per individual, NCBI BioProject ID: PRJNA594029). Reads mapped with a mean success rate of 73.5% to unique locations and 12.9% to multiple locations (Table S4.1). Reads mapped to 37,098 genes, of which 20,387 remained after filtering to remove genes with < 1 cpm in at least 5 individuals in a single treatment-time point combination (Table S4.2). An exploratory PCA highlighted a single individual in the day 80 control treatment as an outlier, which was subsequently removed from downstream analysis (n = 23).

The PERMANOVA for genome-wide gene expression revealed that gene expression differences were associated with treatment (R^2^ = 0.051, *Adonis P* = 0.037) and by time (R^2^ = 0.067, *Adonis P* = < 0.001) but not by their interaction (*Adonis P* = 0.214), indicating a subtle global transcriptome response to both high OA and exposure length (Figure 4A). The DAPC showed that the global patterns of gene expression that differentiated the two treatments at day 9 were not maintained at day 80, with the strongest directional change along the discriminant function in the day 80 samples occurring in the high OA treatment (Figure 4B).

**Figure 4.**
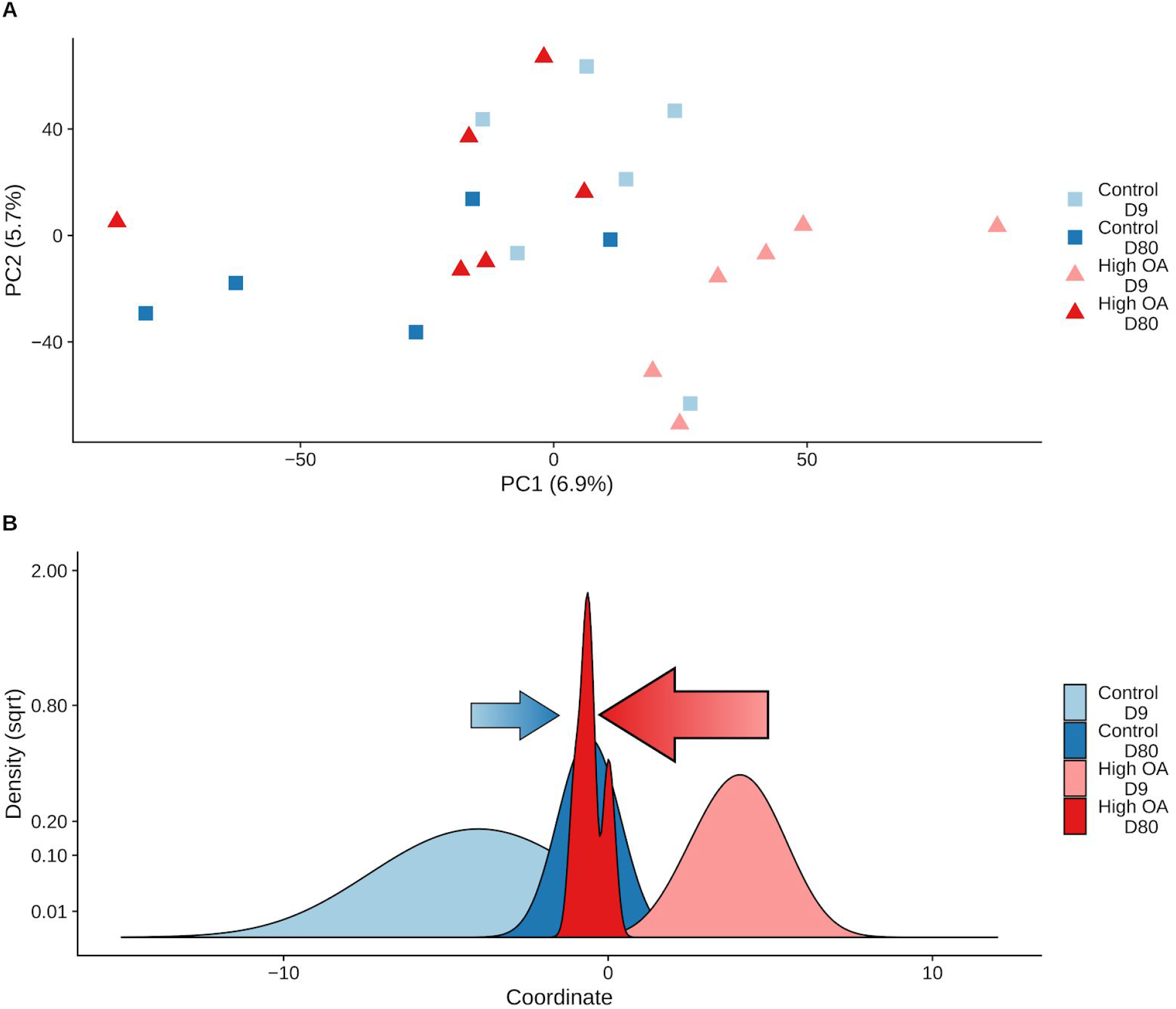
Global transcriptomic responses to OA. **(A)** Plot of the first two principal components from a principal components analysis (P_Trt_ = 0.037, P_Time_ < 0.001, P_TimexTrt_ = 0.214)**. (B)** Density plot of the discriminant values from a DAPC. Discriminant values were determined by using the DAPC package to estimate a function that maximally discriminates between treatment using samples from day 9, then the discriminant value for each sample from day 80 was predicted using the same discriminant function. Arrows indicate direction and degree of movement of expression patterns for control (blue) or OA (red) treatments along the discriminant function from day 9 to day 80. Both plots are based on log2-cpm gene expression.

Interestingly, no genes were differentially expressed. This included a number of genes known from the literature to be associated with biomineralization (Table S4.3). These biomineralization-associated genes tended to be overexpressed relative to the mean expression level for all genes (Figure S4.1), but none were differentially expressed. Furthermore, there was little support to indicate that any of these genes exhibited higher levels of variation in expression among treatment levels (Figure S4.2).

Several GO categories were found to be enriched in genes both up- and down-regulated within the high OA relative to the control treatment (Table 1). Few GO categories appeared to be shared among time points, with the exception of two categories that were enriched in genes up-regulated (‘DNA metabolic process’ and ‘cellular nitrogen metabolic processes’) and one enriched in genes down-regulated (‘transcription factor complex’) in the high OA treatment. Several other metabolic and biosynthetic related biological processes tended to dominate our list of significantly enriched GO categories, suggesting a metabolic response to high OA, particularly on day 80. Up-regulation of several GO categories associated with oxidoreductase activity were also found on day 80.

### Associating DNA methylation with gene expression and EPF pH

All methylomic-transcriptomic comparative analyses (next two sections) were performed using samples with overlapping data (n = 22). Only genes with at least 20% coverage of all CpG were included in the analyses. We argue this represents a reasonable trade-off between maximizing the number of retained genes in the analyses while removing genes with limited or no DNA methylation data (Figure S3.1).

### General DNA methylation and gene expression

Using a PCA approach to investigate the relationship between DNA methylation, gene expression, and other gene attributes, we found the first three PC axes explained >85% of the variance in the data (PC1: 44.8%, PC2: 27%, PC3: 14.9%; Figure 5A-B). PC1 was negatively associated with the length of the gene (*PC axis loading, percent contribution to PC axis*; gene_length: −0.504, 27%), the number of exons (exon: −0.454, 22%), the number of CpGs (totalCpG: −0.484, 25%), and gene expression (Gene_Expression: −0.398, 16%). PC2 was negatively associated with mean methylation (−0.636, 40%), and positively associated with methylation variation among treatments (Methylation_CV: 0.588, 33%). Finally, PC3 was dominated by gene expression (0.482, 23%), variation in expression among individuals (Gene_Expression_CV: −0.672, 46%), and methylation variation among treatments (Methylation_CV: 0.376, 15%). Importantly, these patterns were robust even under increasingly stringent DNA methylation coverage scenarios (Figure S3.2, Figure S3.3). Finally, a follow-up linear regression found that gene expression exhibited a significant positive relationship with methylation (R^2^ = 0.0624, *P* < 0.0001; Figure 5C), while the variation in expression had a significant negative correlation with methylation (R^2^ = 0.0658, *P* < 0.0001; Figure 5D).

**Figure 5.**
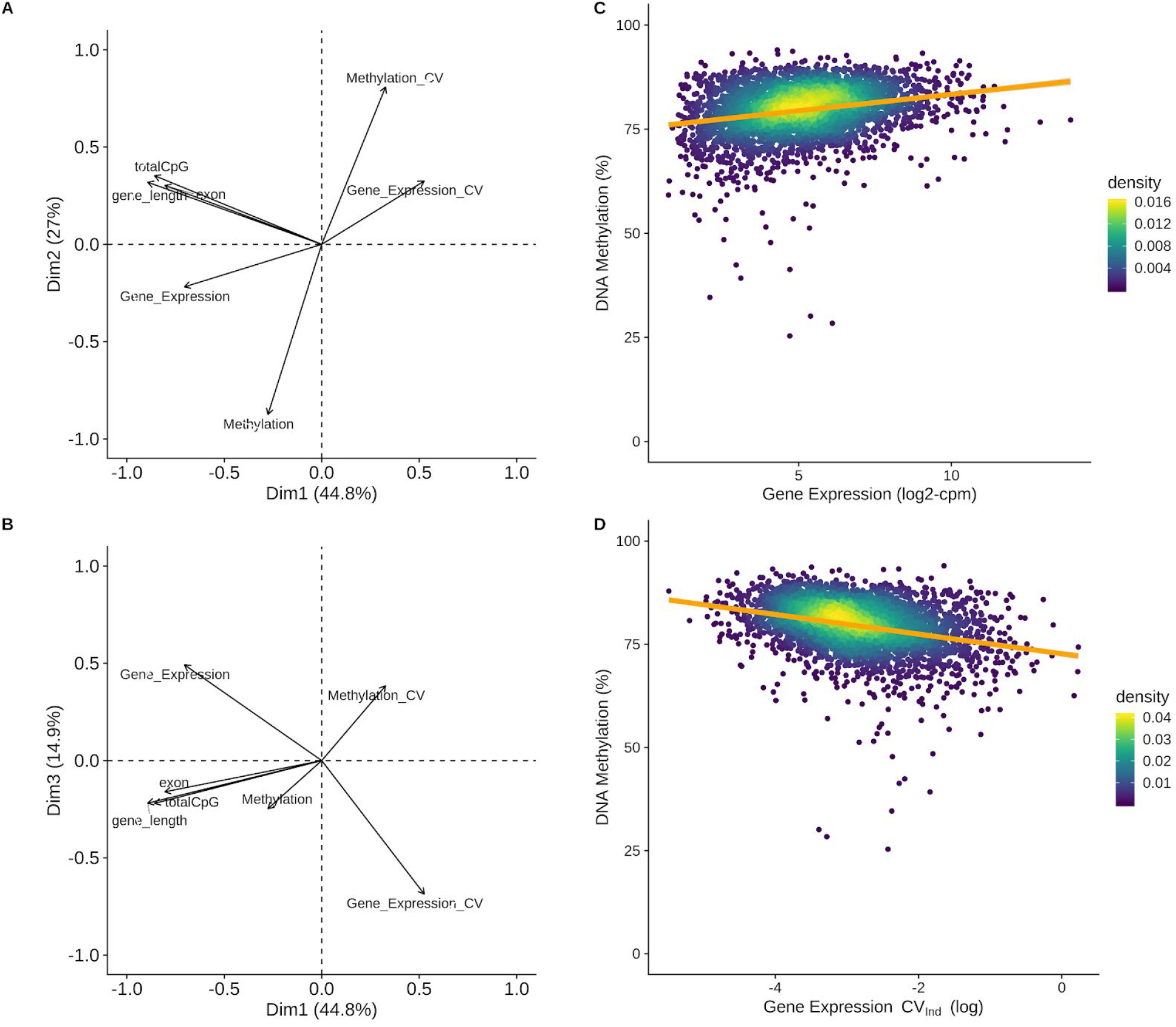
DNA methylation and gene expression correlations. **(A)** First two principal components and **(B)** the first and third components from a principle component analysis that included gene level summary variables for various attributes, expression, and methylation. Variable loadings plotted as arrows, with the length of the arrow corresponding to the relative contribution to PC variance. Significant loadings on the first two PCs included: mean methylation level (Methylation), the coefficient of variance of mean methylation levels among treatments (Methylation_CV), gene expression (Gene_Expression), the number of exons (exon), the number of CpG dinucleotides (totalCpGs), and the gene length in base pairs (gene_length). The CV of gene expression (Expression_CV) was not significant for the first two PCs but was the primary contributor to the third PC (see S3.3). **(C-D)** Plot of gene level DNA methylation against either **(C)** gene expression or the **(D)** gene expression CV among individuals based samples collected in the control treatment at day 9 (Gene Expression - R_2_ = 0.0624, *P* < 0.0001; Gene Expression CV_Ind_ - R_2_ = 0.0658, *P* < 0.0001). Only genes with coverage for at least 20% of all CpGs within the gene were included (n=3,604).

### OA-induced shifts in DNA methylation, gene expression, and EPF pH

A positive relationship was identified via linear regression between the log2-fold change in gene expression and the percent change in mean gene methylation (i.e., average DNA methylation for all CpGs with coverage in a gene; *P_Day9_* < 0.0001, *P_Day80_* < 0.0001, R^2^_Day9_ < 0.012, R^2^_Day80_ = 0.014; Figure 6A), and between the log2-fold change in gene expression and individual DML within the gene among treatments at each time point (*P_Day9_* = 0.0005, *P_Day80_* = 0.0004, R^2^_Day9_ = 0.406, R^2^_Day80_ = 0.276; Figure 6B). Comparing the slopes of these regressions, on average the slopes from the regressions addressing mean gene methylation (slope_Day9_ = 0.0121, slope_Day80_ = 0.0137) were about 25 times greater than the slopes from the regression using DML (slope_Day9_ = 0.0005, slope_Day80_ = 0.00054), indicating that DML had a substantially smaller effect on gene expression.

**Figure 6.**
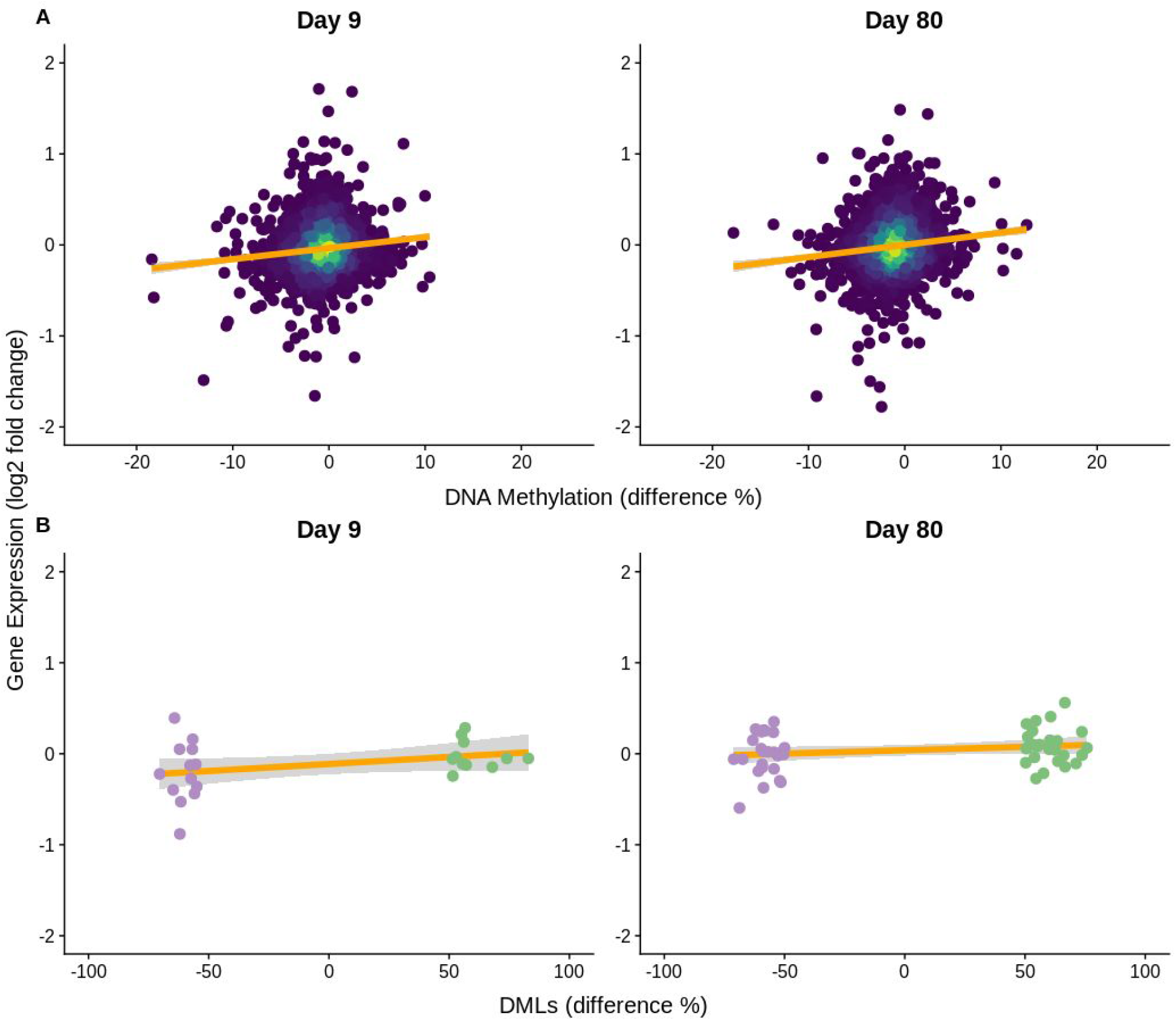
Correlated DNA methylation and gene expression responses to OA. Log2-fold change in gene expression compared to % difference in **(A)** mean gene methylation among treatments (Day 9 - slope = 0.0121, *P* < 0.0001, R^2^ = 0.012; Day 80 - slope = 0.0137, *P* < 0.0001, R^2^ = 0.014) and **(B)** significant DMLs among treatments (Day 9 - slope = 0.0005, *P* = 0.0005, R^2^ = 0.406; Day 80 - slope = 0.00054, *P* = 0.0004, R^2^ = 0.276). Orange line represents the fitted linear model. Colors in **(B)** correspond to DMLs that were significantly hyper- (green) and hypomethylated (purple) in the OA treatment.

Gene co-expression network analysis identified 52 modules of co-expressed genes ranging from 36 to 5059 genes in size (Table S5.1). Of these, the eigengene expression of 8 modules was significantly associated with ΔpH. Next, we examined the potential role of DNA methylation as a mediator of OA in the two modules most significantly associated with ΔpH (cyan and lavenderblush3, which were down-regulated and up-regulated, respectively, as ΔpH increased) and the module most significantly associated with treatment (royalblue, which was up-regulated in the high OA treatment; Figure 7). Note, when selecting the top candidate modules, the module most associated with ΔpH was excluded when the relationship was driven by a single individual outlier. An ANOVA was used to investigate the effect of treatment, time, and their interaction on mean module DNA methylation (calculated at the mean methylation for all CpGs within each gene in the module). Treatment, but not time or their interaction, was significantly associated with the three target modules (*P_Trt,cyan_* = 0.035, *P_Trt,lavenderblush3_* = 0.038, *P_Trt,royalblue_* = 0.033; *P_Time,cyan_* =0.223, *P_Time,lavenderblush3_* =0.557, *P_Time,royalblue_* =0.232; *P_TrtxTime,cyan_* = 0.400, *P_TrtxTime,lavenderblush3_* = 0.202, *P_TrxTime,royalblue_* = 0.671; Figure 7A). Next, a linear regression was used to examine the association between DNA methylation and the eigengene expression for the three target modules, but none were significant (*P_cyan_* = 0.087, *P_lavenderblush3_* = 0.052, *P_royalblue_* = 0.224; Figure 7B). Finally, linear regression was used to look at the correlation between eigengene expression and ΔpH (*P_cyan_* = 0.011, *P_lavenderblush3_* = 0.024, *P_royalblue_* = 0.115; Figure 7C). Both the cyan and lavenderblush3 modules were significantly associated with ΔpH.

**Figure 7.**
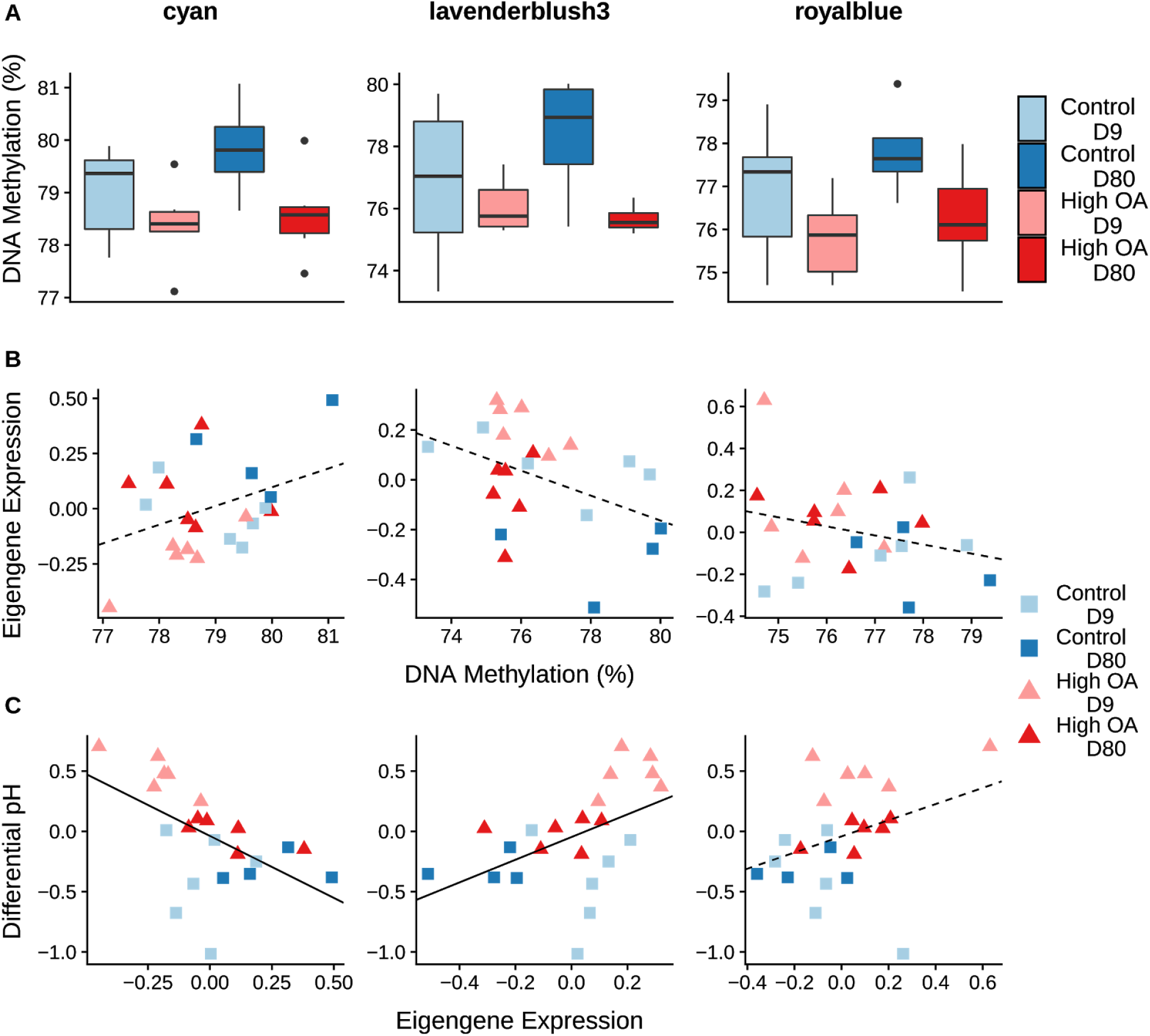
Co-expression and DNA methylation responses to OA. Two of the top three modules associated with ΔpH (cyan and lavenderblush3) and the top module associated with treatment (royalblue). **(A)** Boxplots of mean module methylation (calculated as the mean methylation for all CpGs for each gene within a module) by treatment by day. There was a significant effect of treatment on methylation in all three modules (*P_cyan_* = 0.035, *P_tavenderblush3_* = 0.038, *P_royalblue_* = 0.033), but not a significant effect of time or an interaction. Eigengene expression by **(B)** mean module methylation (*P_cyan_* = 0.087, P_lavenderblush3_ = 0.052, *P_royalblue_* = 0.224) and **(C)** ΔpH (*P_cyan_* = 0.011, *P_lavenderblush3_* = 0.024, *P_royalblue_* = 0.115). Solid lines indicate a significant relationship between explanatory (x-axis) and response variable (y-axis) using a linear model, while dotted lines indicate non significant trends among variables.

## Discussion

In this study we conducted an integrative analysis of physiological, transcriptomic, and methylomic data in the mantle-edge tissue of Eastern oysters to elucidate their molecular response to OA. At the physiological level, oysters tended to maintain a ΔpH significantly lower than pH_seawater_ in our control treatments but not the moderate or high OA treatments, this included day 9 of the exposure where oysters in the the OA treatments maintained ΔpH significantly higher than pH_seawater_. This corresponded with a tendency for the ΔpH in both OA treatments to be significantly higher than the control, consistent with prior work (Liu et al., 2020) showing that eastern oysters regulate their EPF to compensate for acidic conditions. At a molecular level, *C. virginica* exhibited a minor genome-wide response in both the methylome and transcriptome to OA, which partially corresponded with changes in ΔpH. Weak relationships were also observed among DNA methylation and gene expression. Although the ΔpH response was not strongly associated with DNA methylation, significant associations were observed between ΔpH and the eigengene expression of specific gene expression clusters. Below, we discuss how the present study fits into the marine ecological epigenetics and gene expression literature, evaluate whether our data is consistent with the hypothesis that DNA methylation acts as a mediator of gene expression and physiological responses, and explore possible explanations for the subtle patterns in expression and DNA methylation.

### Molecular responses to OA

#### Gene expression

A wide variety of marine calcifying organisms are known to mount significant transcriptomic responses to OA including; corals, urchins, pteropods and other species of oysters (Evans et al., 2013; Davies et al., 2016; Li et al., 2016; Johnson and Hofmann, 2017; Griffiths et al., 2019). Studies of the pearl oyster *Pinctada fuctata*, the purple sea urchin *Strongylocentrotus purpuratus*, and the scleractinian coral *Siderastrea siderea* identified genes associated with ion transport, acid-base regulation, and biomineralization differentially expressed in response to OA (Evans et al., 2013; Evans and Watson-Wynn, 2014; Davies et al., 2016; Li et al., 2016). However, few of these studies have specifically studied the transcriptomic responses within single tissues associated with calcification (but see Hüning et al., 2013; Li et al., 2016). Early transcriptomic work in *C. virginica* showed both genome-wide and individual gene expression differences in the response of gill and hepatopancreas tissue to seawater pH (Chapman et al., 2011), but these tissues are not associated with oyster calcification.

Strikingly, our study found no evidence of OA-induced differential expression of individual genes in the mantle tissue, including a number of biomineralization related genes that were highly expressed relative to other genes in our data. These results are largely consistent with previous work in *C. virginica* mantle tissue, which found limited differences in gene expression in four targeted biomineralization genes in response to OA (Richards, 2017). Notably, despite the lack of differential gene expression in response to OA, they did find significant changes in corresponding protein expression (Richards et al., 2018), suggesting that post-transcriptional regulation is an important mediator of oysters’ phenotypic response to OA. These results contrast a study by Li et al (2016) on *P. fuctata* mantle tissue, which found that approximately 10-11% of genes in their microarray were differentially expressed in response to moderate or severe OA, including a tendency for up-regulation of ion transport genes and down-regulation of biomineralization genes within the severe OA treatment. It is unlikely that the lack of differential gene expression observed in the present study was the result of low statistical power due to small number of biological replicates, as a number of similar studies on marine invertebrates have observed significant gene expression responses to OA with the same number of, or fewer biological replicates (Evans et al., 2013; Li et al., 2016; Johnson and Hofmann, 2017; Griffiths et al., 2019).

Despite the lack of response of individual genes, a broadscale effect of high OA on genome-wide expression was observed. Furthermore, this genome-wide effect changed through time, as evidenced by the enrichment of different GO categories on day 9 compared to day 80 of the OA exposure. Genes that responded to high OA on day 9 (i.e., genes that maximized the difference among OA treatments on day 9 in the DAPC), were not similarly differentiated among treatments on day 80. Moreover, the expression of these genes on day 80 was found to slightly shift toward the expression profile of the control treatment, suggesting that genes responding on day 9 began reverting to control levels of expression by day 80. The GO enrichment analysis showed that both biological processes associated with metabolism and molecular functions associated with oxidative stress were up-regulated in response to OA on day 80, but not on day 9 of the exposure. OA exposure has been shown to elicit oxidative stress responses in oysters, corals, pteropods, and sea urchins (Todgham and Hofmann, 2009; Tomanek et al., 2011; Kaniewska et al., 2012; Wang et al., 2016). A previous study on *C. virginica* mantle tissue also showed that OA exposure is associated with up-regulation of oxygen-stress related enzymes (Tomanek et al., 2011). Similarly, OA exposure in *C. gigas* led to a delayed antioxidant and oxidative stress response in gill tissue, suggesting that redox homeostasis was possible for short term (up to 14 days) but not long term (28 days) exposure to OA (Wang et al., 2016). Transcriptomic studies of corals have found up-regulation of oxidoreductase-related genes in response to OA (Kaniewska et al., 2012), similar to the GO categories identified in this study. In the present study, *C. virginica* exhibited a mild stress response to high OA that is consistent with, but perhaps not as pronounced as, the broad-scale oxidative stress responses observed in other marine calcifiers. Alternatively, the mantle tissue may be constrained or highly polygenic in its response to OA.

#### DNA methylation

*Crassostrea virginica* exhibited a subtle genome-wide DNA methylation response to high OA, similar to the gene expression response of *C. virginica*. This genome-wide DNA methylation response to OA has been found in several other species of marine calcifiers, including corals, pteropods, and in the gonadal tissue of *C. virginica* (Putnam et al., 2016; Liew et al., 2018; Bogan et al., 2020; Venkataraman et al., 2020). In the present study, OA was associated with a genome-wide decrease in methylation (hypomethylation), similar to the response observed in pteropods (Bogan et al., 2020), but opposite the response observed in corals (Putnam et al., 2016; Liew et al., 2018). Over time, an increase in methylation (hypermethylation) was observed, also consistent with the DNA methylation response observed in pteropods (Bogan et al., 2020). Unlike for the pteropods, the same degree of hypermethylation through time was observed in both the control and high OA treatments (i.e., no interaction between treatment and time). Consequently, the hypermethylation that was observed over time could be the result of a minor stress or acclimation response to the experimental setup (consistent with the gene expression results), or a reflection of the temporal instability of DNA methylation in *C. virginica* mantle tissue.

In marine invertebrates, DNA methylation primarily appears to occur in gene bodies (Gavery and Roberts, 2010; Dixon et al., 2018; Zhang et al., 2020), and has been shown to respond to OA (Venkataraman et al., 2020). Consistent with previous findings, the large majority (~94%) of all CpGs with 5x coverage and OA-induced DML were within gene bodies. Interestingly, the overall response of individual loci to OA was quite small (85 DML across both time points) in the mantle tissue compared to the gonadal tissue (Venkataraman et al., 2020), which exhibited hundreds of differentially methylated loci. Although the differential results amongst tissue types may arise from differences in methodologies, the distinct lack of overlap in DML suggests that a fundamental difference in responses exists amongst different tissue types. The 85 OA-induced DML corresponded to a wide range of putative gene functions, including hypomethylation of death-effector domain-containing genes and hypermethylation of a protein-ubiquitination related gene on day 80. Both types of genes are involved in cell apoptosis and possibly reflect a stress response to OA. Conspicuously absent were DML in genes associated with biomineralization-related genes (skeleton development, calcium ion transport, and voltage-gated calcium channel complex), as previously observed in corals (Liew et al., 2018). The lack of overlap in OA-induced DNA methylation in the tissue of performance (mantle) compared to the tissue of inheritance (gonad) in *C. virginica* suggests that parallel responses in the methylome may be limited among tissues, which could have implications for the role of DNA methylation in transgenerational responses of *C. virginica* to OA.

### Linking methylation, expression, and phenotype

Gene body methylation is thought to play a fundamental role in gene regulation across a broad range of taxa (Zilberman et al., 2007; Zemach et al., 2010), and is considered an important mechanism for mediating acclimation response and phenotypic plasticity to stress via gene regulation (Dixon et al., 2018). In marine invertebrates, a positive correlation between gene expression and DNA methylation has been reported in gill and male gametes in *C. gigas* (Gavery and Roberts, 2013; Olson and Roberts, 2014), mantle tissue in *P.fuctata* (Zhang et al., 2020), and across the entire organism in the coral *S.pistillata* (Liew et al.,2018). In response to OA specifically, Liew et al. (2018) found that significant gene expression and calcification corresponded with correlated shifts in DNA methylation, suggesting that DNA methylation may play a role in mediating coral response to OA. Consistent with these previous findings, gene body DNA methylation in *C. virginica* mantle tissue exhibited a subtle but significant association with both gene expression (positive) and gene expression variation (negative), indicating gene body hypermethylation may drive both an increase in gene expression and a decrease in transcriptional variation. However, the small R^2^ and slope from the regressions do not provide strong evidence that gene body DNA methylation had a substantial effect on gene regulation in *C. virginica*. Moreover, no overlap was observed between GO categories enriched in differentially expressed genes and those enriched in differentially methylated loci, nor was there evidence that gene body DNA methylation had a significant effect on the expression of co-expressed gene modules that were significantly associated with one of the primary physiological responses of the oysters (ΔpH).

### Subtle shifts and constraints

The subtle molecular responses of *C. virginica* found in this study contrasted the more severe transcriptomic responses exhibited by other marine calcifying taxa in response to OA (e.g., Li et al. 2016). However, other examples of subtle gene expression shifts in response to environmental change do exist (Malik et al., 2019), and these minor shifts can still have phenotypic consequences (Wu et al., 2010). In *C. virginica*, these subtle molecular responses may be the result of; (i) extended evolution under regular extreme OA conditions, (ii) constrained plasticity of the mantle (e.g., an evolved canalization of gene expression in response to OA), and/or (iii) a highly polygenic genetic response architecture.

*Crassostrea virginica* inhabits coastal estuaries, which can experience regular, unpredictable, and rapid swings in acidity leading to conditions near or above the high OA scenario (see S1.1). Since the high OA treatment is not unusual from the oyster’s perspective, the subtle molecular responses may reflect that the prescribed OA conditions are not sufficiently different from what the oysters’ typically experience on diurnal and/or seasonal cycles so as to elicit a strong molecular response. On the other hand, highly uncertain environments, in which the frequency of fluctuations is much higher than what the organism can respond to or is able to predict, can lead to the evolution of a single expressed phenotype (in this case, gene expression in the mantle) across all environments that maximizes the geometric mean fitness (Seger and Brockmann, 1987; Starrfelt and Kokko, 2012; Botero et al., 2015). The variable environments inhabited by the oysters in the present study may consequently select for constrained (e.g., canalized) responses in the transcriptome and methylome, in which the expression profile results from the evolution of a nearly fixed bet-hedging strategy, rendering the genome capable of only subtle shifts. From an energetic cost perspective, one would expect selective pressure against increases in mRNA and protein expression due to the costs associated with increased expression (Wagner, 2005; Weiße et al., 2015). Interestingly, calcification genes were associated with higher expression than the genome-wide background in the mantle tissue, suggesting that their expression may be maximized, which could constrain their ability to be further up-regulated in response to OA.

Alternatively, subtle shifts may be observed if the traits responding to OA are highly polygenic (Pritchard et al., 2010) and/or there is a redundant genetic architecture (Yeaman, 2015). Although this body of literature is focused primarily on shifts in allele frequencies in response to selection rather than gene expression, the same principles about polygenic traits apply. In the case of plastic polygenic traits, the magnitude of response of individual genes may be relatively small and, therefore, not detectable via differential expression analysis. In the case of redundant genetic architecture, theory predicts that high genetic redundancy in a population occurs when individuals can reach a phenotypic outcome by a wide range of genes and pathways (see topic reviewed in Láruson et al., 2020), which could result in very large residual error and limited statistical power to detect differences amongst treatment (i.e., lack of a clear significant response because every individual does something different to achieve the same phenotype). Regardless of whether these results are more influenced by subtle shifts or constraints on gene expression, they show that a marine calcifier’s drastic phenotypic responses to OA (i.e., pH_EPF_ response and reduced calcification rate under high OA) can be associated with only subtle molecular responses within that organism’s calcifying tissue.

## Conclusion

*Crassostrea virginica* exhibited reduced calcification, reduced pH_EPF_, and increased ΔpH in response to long-term (80-day) exposure to CO_2_-induced ocean acidification. These responses to OA were relatively stable throughout the 80-day experiment, although elevation of ΔpH under the high OA treatment was enhanced at day 9 and pH_EPF_ was highly variable within all treatments during the first 22 days of the experiment. The observed increase in ΔpH under the moderate and high OA conditions may support calcification within *C. virginica* during ephemeral acidification events, such as those they already experience due to diurnal and seasonal fluctuations in seawater pH. Notably, we did not find a strong association between DNA methylation and OA-induced gene expression or ΔpH. This was likely due, in part, to the surprisingly subtle responses to OA in both the methylome and transcriptome of the mantle tissue. Since the majority of OA gene expression studies have not specifically targeted calcifying tissue, our results highlight a need to re-assess how flexible the oyster calcification process is at the molecular level. Additional studies targeting the specific molecular response of calcifying tissues are needed to better understand how organisms are able to regulate calcification processes in response to OA.

## Supporting information

General Supplemental

## Data Availability

All scripts and data will be uploaded to the BCO-DMO database after the paper is accepted but before publication.

**Current location of scripts, data, and pipeline** https://github.com/epigeneticstoocean/AE17_Cvirginica_MolecularResponse

**BioProject ID:** PRJNA594029

## Acknowledgements

We acknowledge Sara Schaal, Aki Laruson, Isaac Westfield and other members of the Lotterhos and Ries labs for their helpful comments that improved the manuscript. This research was funded by the National Science Foundation (1635423). Resources purchased with funds from the NSF FMSL program (DBI 1722553, to Northeastern University) were used to generate data for this manuscript. We are also thankful for the help of many graduate students, undergraduates, technicians, and volunteers during the experiment, including Andrea Unzueta Martinez, Bodie Weedop, Isabel Gutowski, Jaxine Wolfe, Sandi Scripa, Kevin Wasko, Isaac Westfield, and Camila Cortina.

## Author Contributions Statement

Principle design of the experiment was done by ADW, SR, JR, and KL. Field collections were done by ADW, BF, and EM. The experiment was conducted by ADW. During the experiment, extrapallial fluid was collected and analyzed by LC, while other phenotypic data and tissues were collected by ADW. Water chemistry was analyzed by ADW and EM. Tissues were prepared for RNAseq sequencing by BF and prepared for MBD-BSseq by YV and SR. All bioinformatics and statistical analysis was performed by ADW. The manuscript was written by ADW with assistance from KL, LC, JR, and YV. All authors contributed to editing and approving the final version of the manuscript.

