## Supplementary material for "Ocean acidification induces subtle shifts in gene expression and DNA methylation in mantle tissue of the Eastern oyster (*Crassostrea virginica*)": General Supplemental

### Supplementary Data

#### Table of Contents


**Supplemental Section 1: Water Chemistry and Field Data** **2**

Table S1.1 Experimental water chemistry summary 3

Figure S1.1 Field site water chemistry. 5

Figure S1.2 Mean triweekly water chemistry. 6

**Supplemental Section 2: Phenotype Data** **7**

Figure S2.1 Buoyant weight compared to dry weight. 8

**Supplemental Section 3: DNA Methylation Data** **9**

Table S3.1 DNA Methylation mapping summary. 10

Table S3.2 DNA Methylation CpG by genomic feature summary. 11

Table S3.3 Differentially methylated CpGs (DMLs) among treatments. 12

Figure S3.1 Gene by CpG Coverage 13

Figure S3.2 Gene Summary PCAs 14

Figure S3.3 Principle Component Contributions 16

**Supplemental Section 4: Transcriptomic Data** **18**

Table S4.1 Per sample read and mapping summary for RNAseq data. 19

Table S4.2 Gene expression quantification summary. 20

Table S4.3 Target list of genes associated with biomineralization in marine calcifiers from the literature. 21

Figure S4.1 Density plot of expression in all genes vs. biomineralization target list 22

Figure S4.2 Expression variation in all genes vs. biomineralization target among treatment levels 23

**Supplemental Section 5: Comparative** **24**

Table S5.1 WGCNA Module Summary 24

### Supplemental Section 1: Water Chemistry and Field Data

### Table S1.1 Experimental water chemistry summary

Measured water chemistry including temperature (C$^o$), salinity (PSU), and pH (NBS scale) were measured Monday, Wednesday, and Friday. Dissolved carbon (DIC) and total alkalinity (AT) were measured every two weeks. Calculated water chemistry including, calculated pH (SW scale), $pCO_2$, bicarbonate ion concentration ($HCO_3^-$), carbonate ion concentration ($CO_3^{2-}$), dissolved $CO_2$ ($CO_{2,SW}$), calcite saturation state ($\Omega$ Ca), and atmospheric $pCO_2$. Note, whenever possible DIC and AT were used to calculate the complete carbonate chemistry, but measured pH was substituted in when DIC or AT was not available.

| Measured Parameter | | Acc. | Exposure Period | | | | | |
|---|---|---|---|---|---|---|---|---|
| | | | Day 9 | | | Day 80 | | |
| | | | Control | OA 1000 | OA 2800 | Control | OA 1000 | OA 2800 |
| N (C,psu,pH) | | 198 | 36 | 36 | 36 | 186 | 186 | 186 |
| **Temp.** | **(celsius)** | 17.0 | 17.4 | 17.3 | 17.3 | 18.3 | 18.4 | 18.4 |
| | **SEM** | 0.03 | 0.08 | 0.08 | 0.08 | 0.08 | 0.08 | 0.08 |
| **Salinity** | **(psu)** | 29.91 | 30.27 | 30.28 | 30.28 | 31.03 | 31.06 | 31.06 |
| | **SEM** | 0.03 | 0.03 | 0.03 | 0.03 | 0.03 | 0.03 | 0.03 |
| **pH** | **(NBS)** | 7.77 | 7.82 | 7.55 | 7.14 | 7.83 | 7.57 | 7.10 |
| | **SEM** | 0.00 | 0.01 | 0.01 | 0.01 | 0.01 | 0.01 | 0.01 |
| **N (DIC and AT)** | | 23 | 12 | 9 | 9 | 32 | 32 | 35 |
| **DIC** | **(µmol/kg)** | 1904.22 | 1901.92 | 1991.29 | 2161.02 | 1989.83 | 2066.89 | 2215.22 |
| | **SEM** | 3.06 | 5.69 | 12.62 | 11.15 | 3.53 | 3.97 | 11.22 |
| **AT** | **(µmol/kg)** | 2043.58 | 2050.38 | 2069.06 | 2101.63 | 2146.79 | 2142.99 | 2175.36 |
| | **SEM** | 4.43 | 7.20 | 5.59 | 11.04 | 1.69 | 3.79 | 4.11 |
| **Calculated Parameters** | | | | | | | | |
| **N** | | 25 | 12 | 12 | 12 | 32 | 32 | 36 |
| **pH calc** | **(SW scale)** | 7.90 | 7.90 | 7.72 | 7.26 | 7.89 | 7.66 | 7.29 |
| | **SEM** | 0.01 | 0.02 | 0.03 | 0.02 | 0.01 | 0.02 | 0.03 |
| **pCO2** | **(uatm)** | 547.3 | 546.0 | 879.8 | 2705.7 | 579.1 | 1050.4 | 2728.6 |
| | **SEM** | 20.2 | 22.3 | 57.8 | 107.2 | 16.5 | 47.5 | 128.0 |
| **HCO3-** | **(µmol/kg)** | 1771.2 | 1768.0 | 1866.4 | 2017.5 | 1847.5 | 1952.9 | 2083.2 |
| | **SEM** | 4.7 | 6.9 | 13.4 | 10.5 | 4.8 | 4.7 | 10.1 |
| **CO32-** | **(µmol/kg)** | 111.0 | 114.6 | 80.1 | 30.0 | 122.2 | 77.6 | 37.0 |
| | **SEM** | 2.9 | 3.7 | 3.7 | 1.0 | 2.0 | 2.2 | 3.2 |
| **CO2 (SW)** | **(µmol/kg)** | 19.83 | 19.34 | 31.28 | 95.93 | 20.17 | 36.36 | 94.85 |
| | **SEM** | 0.72 | 0.80 | 1.97 | 3.71 | 0.57 | 1.57 | 4.30 |
| **Omega Ca** | | 2.74 | 2.82 | 1.97 | 0.74 | 3.00 | 1.91 | 0.91 |
| | **SEM** | 0.07 | 0.09 | 0.09 | 0.02 | 0.05 | 0.05 | 0.08 |
| **pCO2 (gas)** | **(ppm)** | 557.8 | 556.9 | 897.2 | 2759.7 | 591.0 | 1072.4 | 2785.0 |
| | **SEM** | 20.6 | 22.7 | 59.0 | 109.5 | 16.9 | 48.6 | 130.9 |

Figure S1.1 Field site water chemistry.

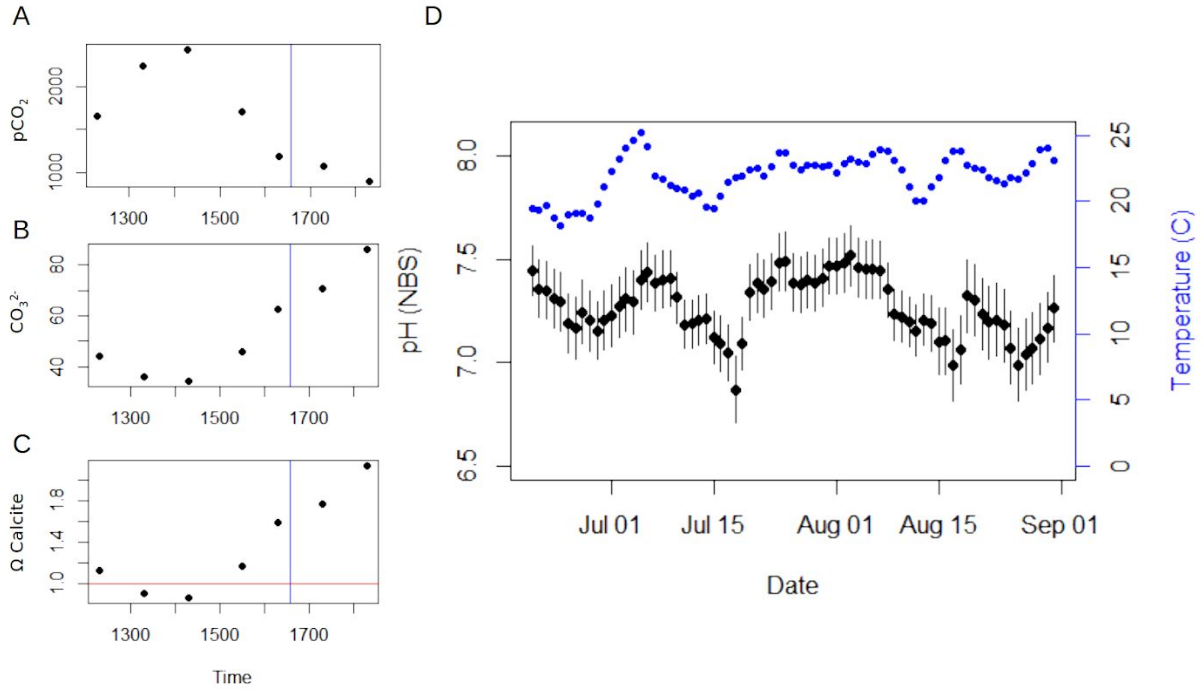


**Figure S1.1 | Field site water chemistry.** Carbonate chemistry and temperature from collection site 3 (42.681764, -70.813498) including a detailed tidal cycle measurement (A-C) and daily average pH and temperature summary (D). In summer 2016, seven samples were collected for alkalinity and DIC starting approximately 3 hours prior and continuing until 3 hours post low tide. Blue lines represent the low tide for the nearest NOAA buoy, which had an approximately 2 hour lag compared to the collection site. Measured salinity, temperature, alkalinity and DIC were used in $CO_2$Sys to determine the complete carbonate chemistry including, $pCO_2$ (A), carbonate (B), and calcite saturation state (C). The red line indicates the critical calcite saturation state. In panel D, daily mean pH (black diamonds, mean±SE) and temperature (blue circles) over the course of summer 2017.

### Figure S1.2 Mean triweekly water chemistry.

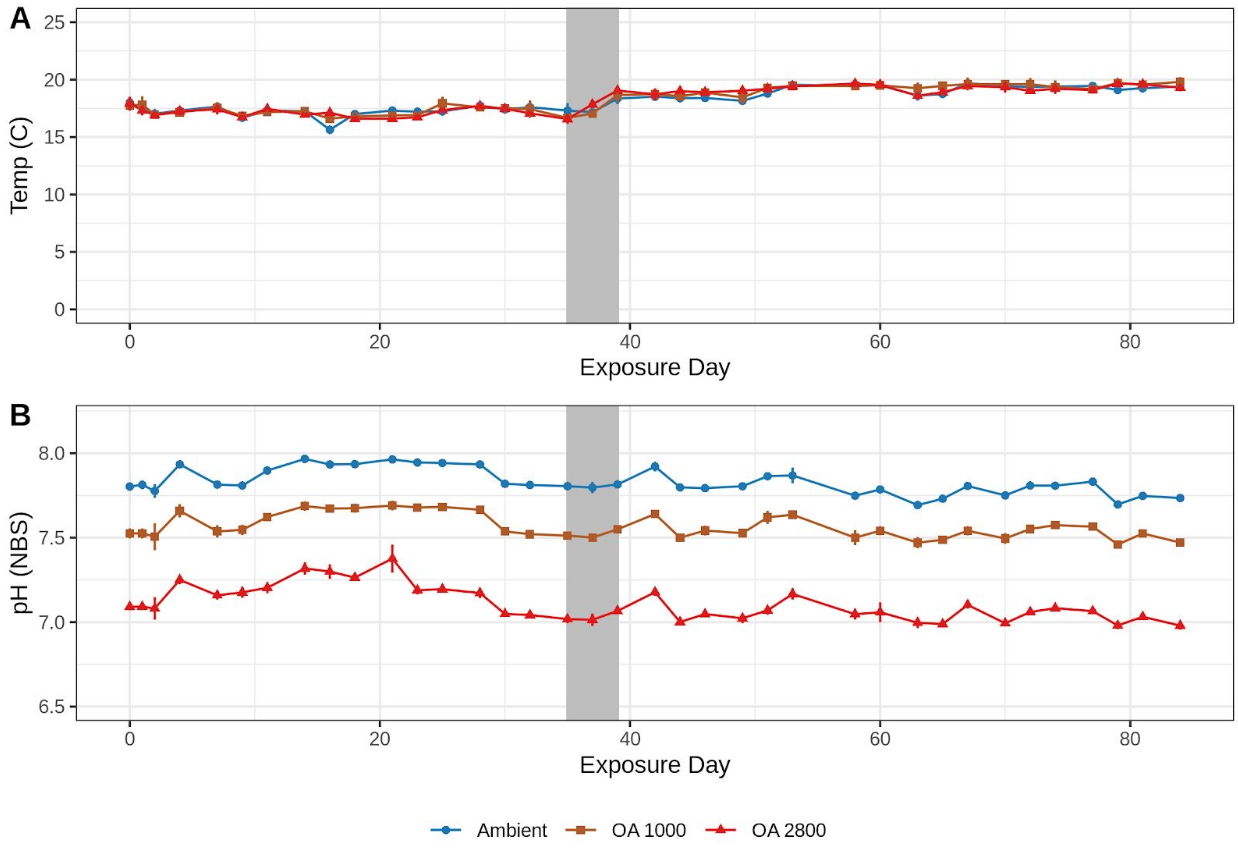


**Figure S1.2 | Mean triweekly water chemistry.** Mean temperature **(A)** and pH **(B)** during the experimental exposure. Lines represent means across the 6 replicate tanks per treatment with vertical bars showing the 95% CI. Grey box highlights the ~5 day period where the temperature was increased by 1.5 degrees C.

### Supplemental Section 2: Phenotype Data

Figure S2.1 Buoyant weight compared to dry weight.

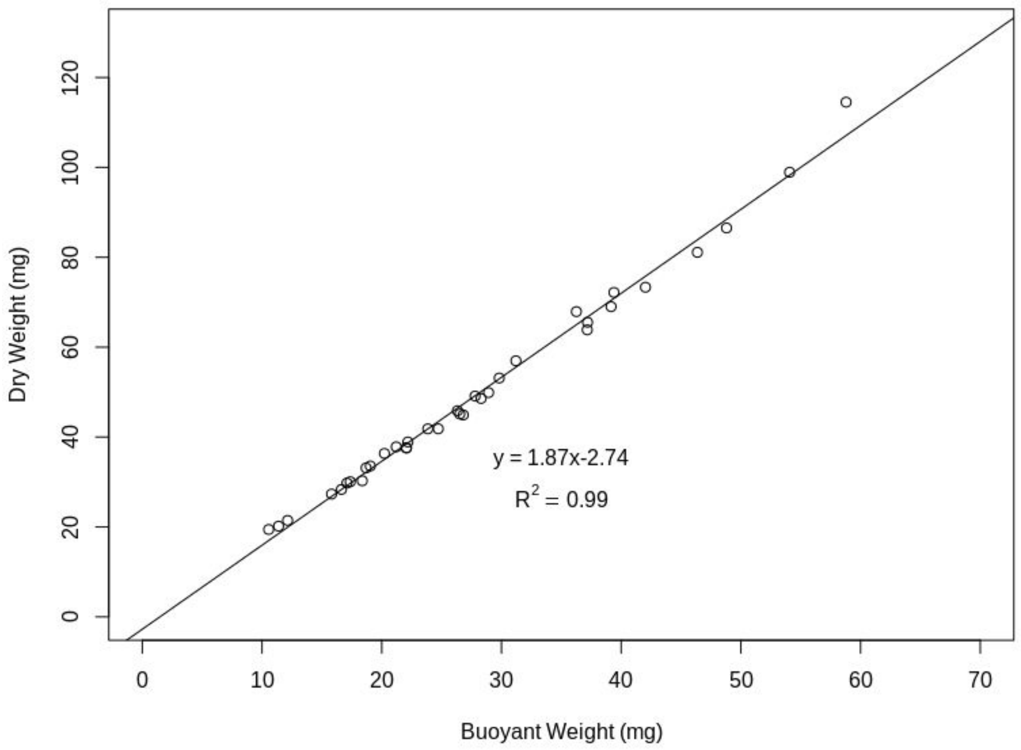


**Figure S2.1 | Buoyant weight compared to dry weight.** Buoyant and dry weights for oysters from oysters collected during the first and second buoyant weight measuring timepoints. Linear regression was used to evaluate the relationship between the two variables, estimate the slope and intercept, and determine the degree of correlation.

### Supplemental Section 3: DNA Methylation Data

### Table S3.1 DNA Methylation mapping summary.

Trimming and quality control performed using Trim_Galore! with automated adapter detection and clipping 10bp from both 5’ and 3’ ends of each read. Mapping was performed in bismark with the bowtie2 mapper using a score_min setting of L0,0,-0.8.

| **Step** | **Total** | **Per Sample** | | | |
|---|---|---|---|---|---|
| | | **Mean** | **SD** | **Min** | **Max** |
| Sequencing Reads (millions) | 1409.8 | 59.99 | 8.43 | 37.6 | 71.6 |
| Reads Mapped (millions) | 622.4 | 25.93 | 3.88 | 17.1 | 31.9 |
| Reads Mapped (percent) | NA | 43.35% | 0.02% | 39.40% | 46.00% |
| Reads Mapped After deduplication (millions) | 566.3 | 23.60 | 4.41 | 10.2 | 28.7 |

#### Table S3.2 DNA Methylation CpG by genomic feature summary.

Counts based on an estimate of the total number of CpGs in the oyster genome (Accession: GCA_002022765.4). The threshold represents minimum per sample coverage (>1, >=5, >=10) for CpG inclusion. Based on 23 samples after removing one individual due to poor sequencing.

| | **All CpG Destranded** | **Sequenced CpG covered (>=1x)** | **Sequenced CpGs (>=1x per sample)** | **Sequenced CpGs (>=5x per sample)** | **Sequenced CpGs (>=10x per sample)** |
|---|---|---|---|---|---|
| Num. of CpGs | 14,458,703 | 12,765,452 | 932,973 | 403,976 | 294,911 |

#### Table S3.3 Differentially methylated CpGs (DMLs) among treatments.

Summary table of differentially methylated CpGs using the logistic regression approach in methylKit. These include the 2 loci where treatment had a significant effect and the 83 loci which were significant by treatment on either day 9 or 80. Attached as a separate excel file.

### Figure S3.1 Gene by CpG coverage

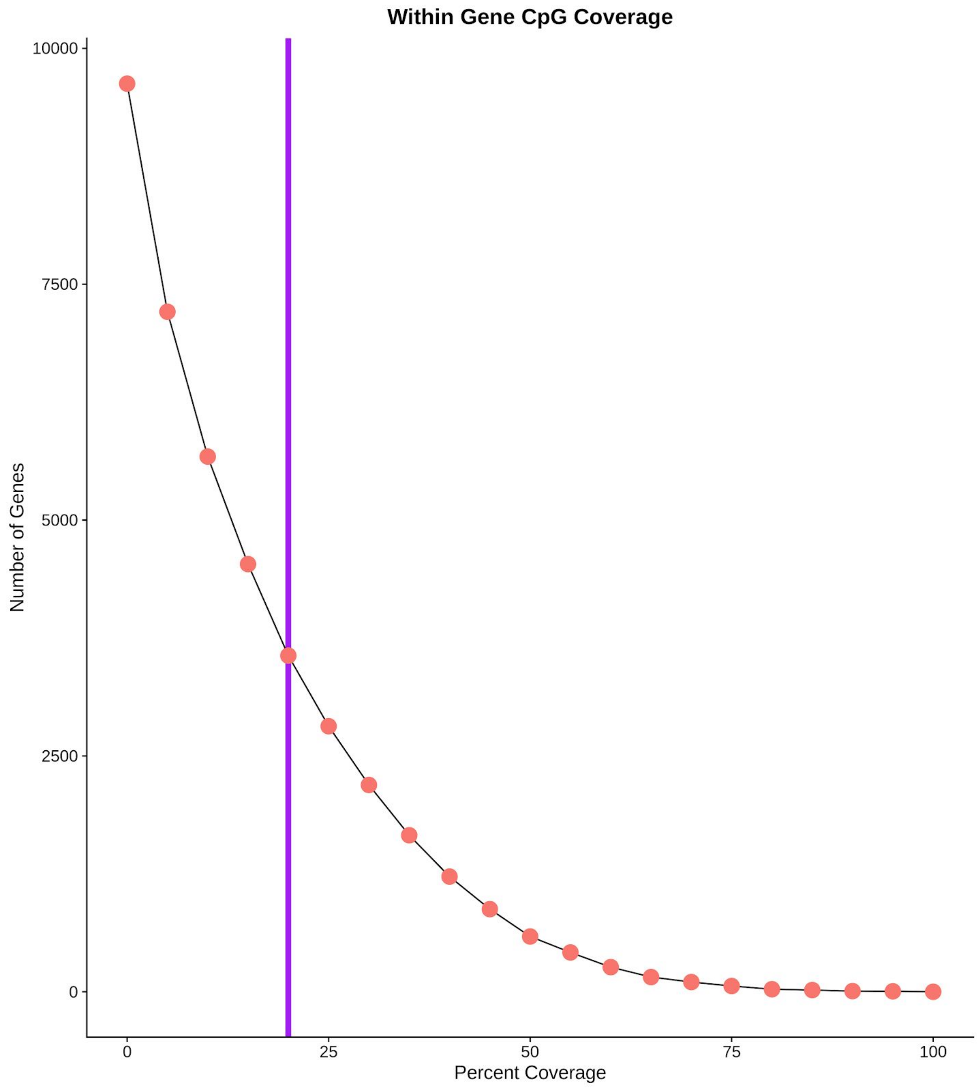


**Figure S3.1 | Gene by CpG Coverage.** Orange circles show the number of genes that have a minimum percent of CpGs covered within a gene for a range of percent thresholds. The purple line indicates the 20% minimum coverage threshold used.

### Figure S3.2 Gene summary PCAs

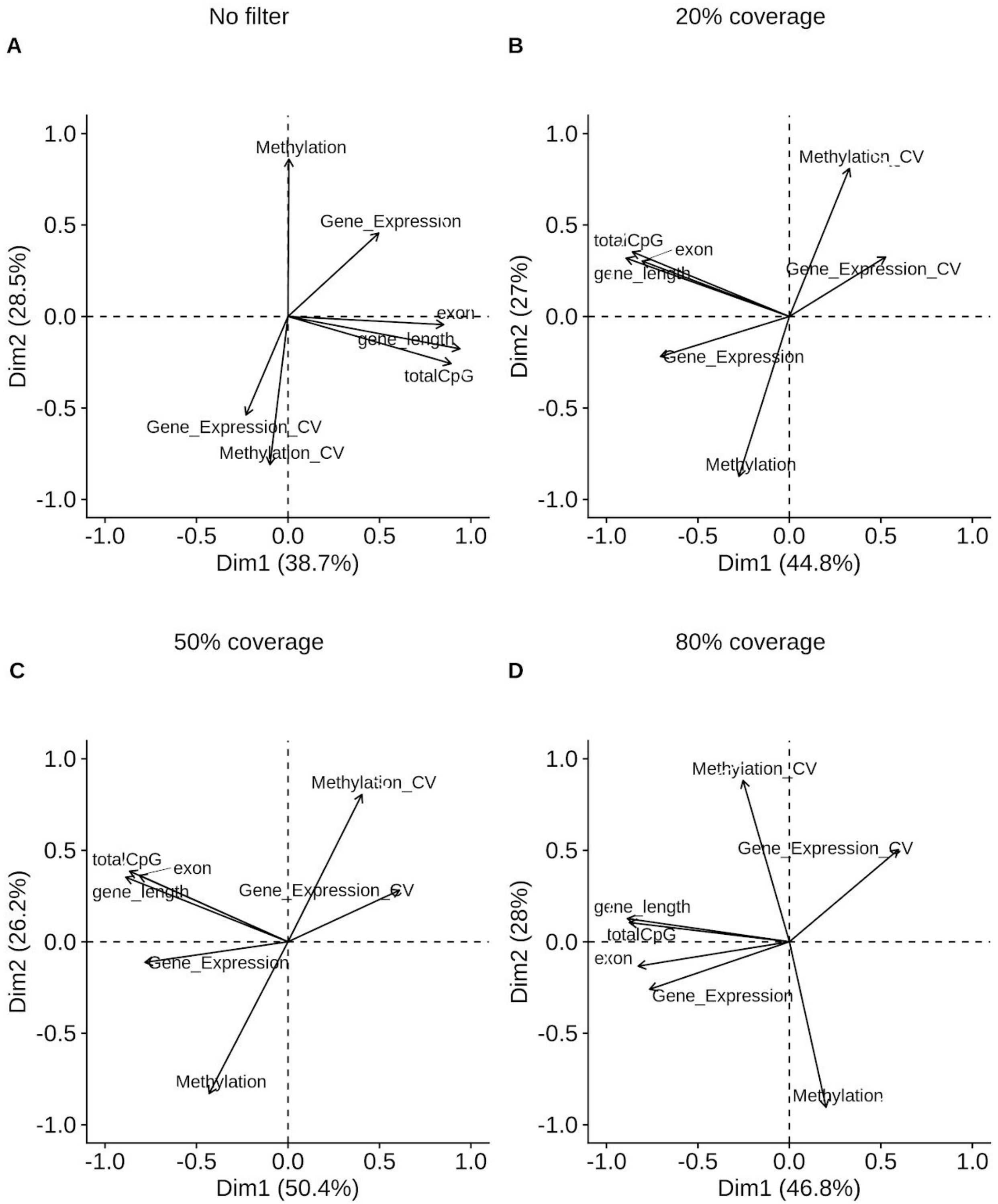

**Figure S3.2 | Gene Summary PCAs for all minimum thresholds.** First two principal components from a principle component analysis that included gene level summary variables for various attributes, expression, and methylation based on **(A)** no filter, **(B)** 20%, **(C)** 50%, **(D)** and 80% coverage thresholds. Thresholds based on the percent of CpGs with the minimum sequence coverage (i.e., 20% coverage only includes genes with at least 20% CpG sequence coverage in the PCA).Variable loadings plotted as arrows, with the length of the arrow corresponding to the relative contribution to PC variance.

### Figure S3.3 Principle component contributions

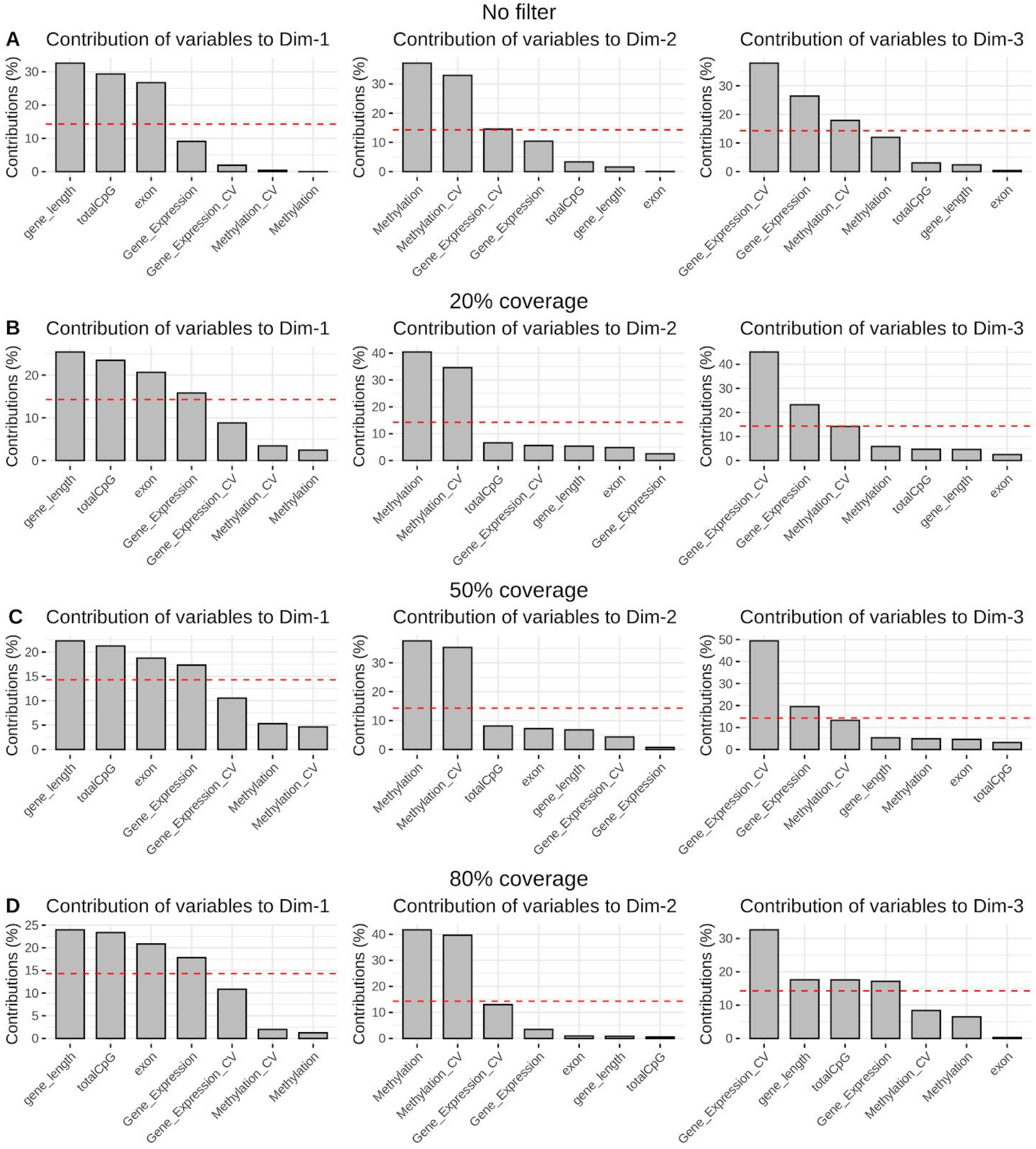

**Figure S3.3 | Principle component contributions.** Relative contribution (percent) for each variables (loading) for the first (column 1), second (column 2), and third (column3) principal components of the PCA for all genes **(A)**, genes with coverage for at least 20% **(B)**, 50% **(C)**, and 80% **(D)** of all CpGs. The dotted redlines indicate the default 15% contribution significance threshold for individual variables.

### Supplemental Section 4: Transcriptomic Data

### Table S4.1 Per sample read and mapping summary for RNAseq data.

Trimming and quality control performed using a custom pipeline implemented in dDocent and using trimmomatic. Mapping performed using the aligner STAR with the 2pass procedure. Statistics are calculated over samples (*n* = 24).

| | MEAN | SD | MIN | MAX |
|---|---|---|---|---|
| Sequencing Reads (per sample) | 39,750,000 | 2,670,000 | 36,150,000 | 45,050,000 |
| Sequencer Read Quality | 39.0 | 0.1 | 38.8 | 39.1 |
| Number Read After Trimming and QC | 27,930,000 | 2,260,000 | 24,390,000 | 32,740,000 |
| STAR - Unique_reads | 20,540,000 | 1,770,000 | 18,200,000 | 24,630,000 |
| STAR - Unique_Percent | 73.5% | 1.4% | 69.0% | 75.3% |
| STAR - Multi_reads | 3,600,000 | 280,000 | 3,130,000 | 4,130,000 |
| STAR - Multi_Percent | 12.9% | 0.8% | 12.0% | 14.8% |

### Table S4.2 Gene expression quantification summary.

Mean gene expression counts and number of putative genes before and after filtering (n = 24). Filtering included removing genes that did not contain at least 1 transcript per million in at least 5 (out of 6) samples in at least one treatment and time level.

| **Gene Quantification Method** | **Mean** | **SD** | **Min** | **Max** | **Number of genes** |
|---|---|---|---|---|---|
| Total | 13,456,348 | 1,271,404 | 11,592,273 | 16,427,649 | 37,098 |
| After Filtering | 13,320,344 | 1,256,664 | 11,474,534 | 16,257,768 | 20,387 |

### Table S4.3 Target list of genes associated with biomineralization in marine calcifiers from the literature.

**(A)** Table of biomineralization genes from the literature and their expression within our data. **(B)** Full summaries of all gene names and locations in reference to the oyster genome (NCBI BioProject ID: PRJNA594029) and **(C)** summary of literature sources. Attached as a separate excel file.

Figure S4.1 Density plot of expression in all genes vs. biomineralization target list

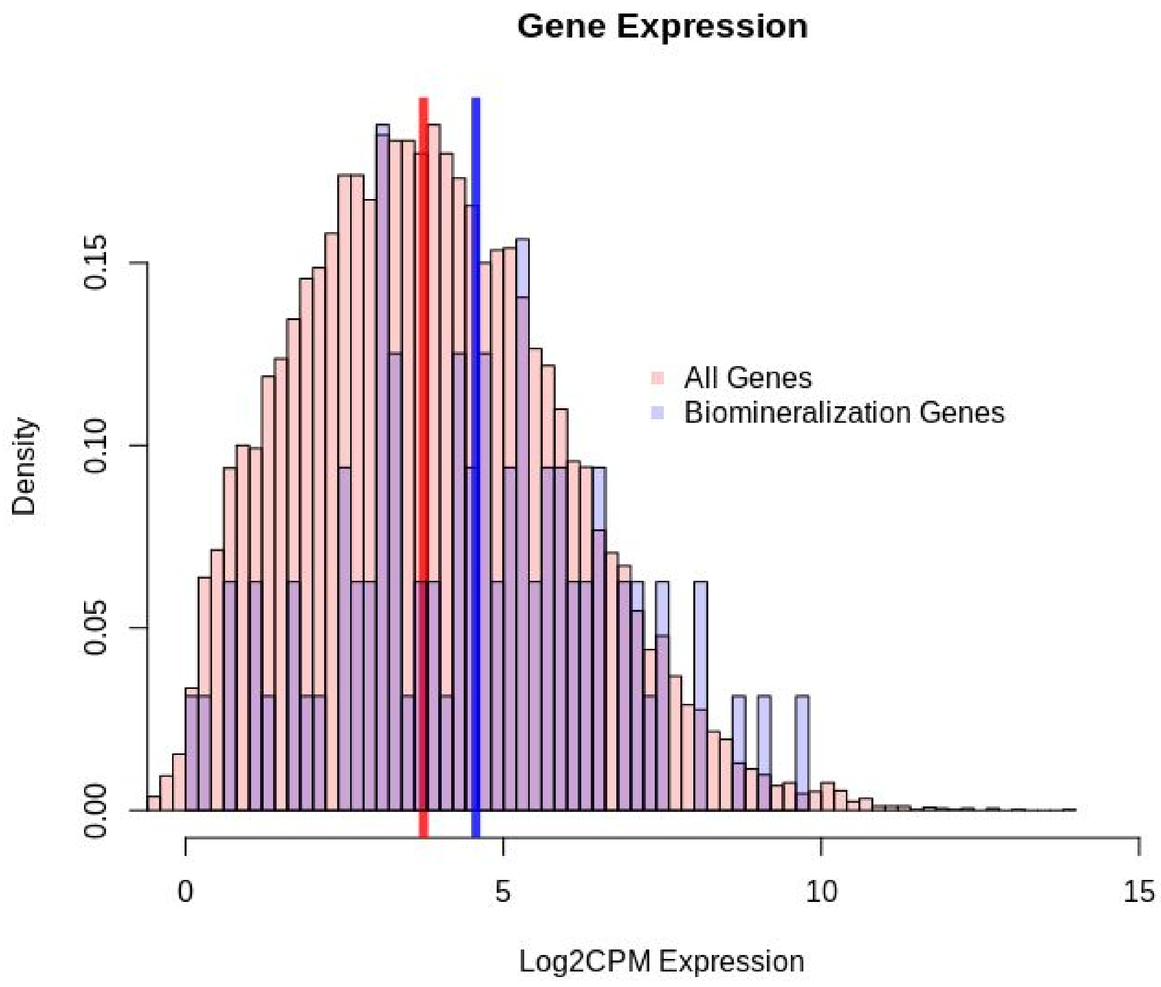


**Figure S4.1 | Density plot of expression in all genes vs. biomineralization genes.** Dark vertical lines indicate the median expression for all genes (red, n = 20387) and biomineralization genes (blue, n = 90); these represent statistically different medians (*P* = 0.002, Mann Whitney U rank sum).

Figure S4.2 Coefficient of variation in gene expression of all genes vs. biomineralization target list

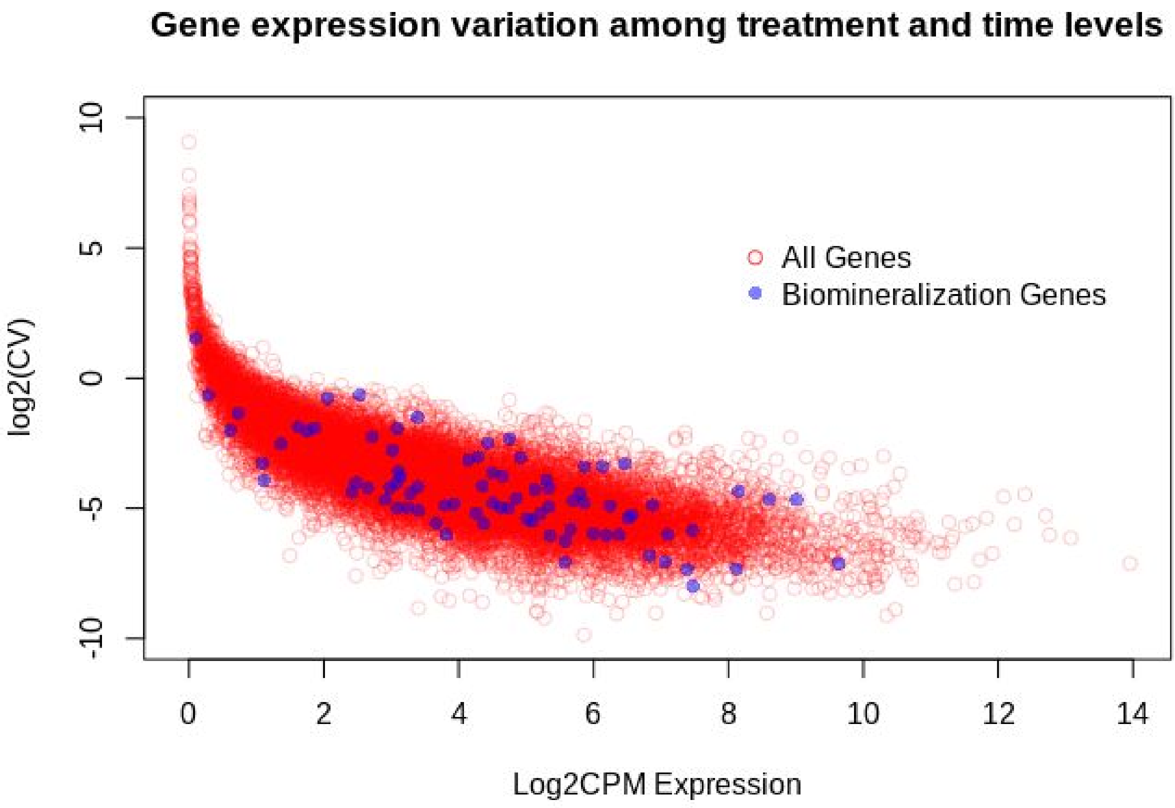


**Figure S4.2 | Coefficient of variation in gene expression of all genes vs. biomineralization target list.** Coefficient of variation (CV) was calculated based on the mean log2-cpm expression among all treatment and time combinations for all genes (red, n = 20387) and the target biomineralization genes (blue, n = 90).

### Supplemental Section 5: Comparative

#### Table S5.1 WGCNA module summary

**(A)** Summary of the 52 different co-expression gene modules generated with WGCNA, including number of genes in the module, eigenexpression, percent methylation, proportion of CpG with coverage, and model statistics (P and R2 values) for each comparison. Values in bold indicate significant effects. The asterisk (*) indicate the modules used for Figure 7. (**B-D**) Gene list summaries of the three top module candidates. Attached as a separate excel file.
